# Antiviral defense in aged *Caenorhabditis elegans* declines due to loss of DRH-1/RIG-I deSUMOylation via ULP-4/SENP7

**DOI:** 10.1101/2024.11.12.623310

**Authors:** Yun Zhang, Andrew V. Samuelson

## Abstract

Innate host defense mechanisms require posttranslational modifications (PTM) to protect against viral infection. Age-associated immunosenescence results in increased pathogenesis and mortality in the elderly, but the contribution of altered PTM regulation to immunosenescence is unknown. SUMOylation is a rapid and reversible post-translational modification that has been implicated in age-associated disease and plays conflicting roles in viral replication and antiviral defenses in mammals. We have discovered in *Caenorhabditis elegans* that induction of antiviral defense is regulated through SUMOylation of DRH-1, the ortholog of the DEAD/H-box helicase and cytosolic pattern recognition receptor RIG-I, and that this regulation breaks down during aging. We find the SUMO isopeptidase ULP-4 is essential for deSUMOylation of DRH-1 and activation of the intracellular pathogen response (IPR) after exposure to Orsay virus (OV), a natural enteric *C. elegans* pathogen. ULP-4 promotes stabilization of DRH-1, which translocates to the mitochondria to activate the IPR in young animals exposed to virus. Loss of either *drh-1* or *ulp-4* compromises antiviral defense resulting in a failure to clear the virus and signs of intestinal pathogenesis. During aging, expression of *ulp-4* decreases, which results in increased proteosomal degradation of DRH-1 and loss of the IPR. Mutating the DRH-1 SUMOylated lysines resulted in the constitutive activation of the IPR in young animals and partially rescued the age-associated lost inducibility of the IPR. Our work establishes that aging results in dysregulated SUMOylation and loss of DRH-1, which compromises antiviral defense and creates a physiological shift to favor chronic pathological infection in older animals.

## Introduction

Viral infection is universal to all organisms. However, the physiological state of the host plays a major role in shaping host-viral interactions and the outcome of an infection. Numerous factors influencing an organism’s physiological status: environmental conditions, diet and nutrition, sleep, comorbidities, genetic polymorphisms, and aging can all influence the severity and outcome of an acute infection (1). For instance, the COVID-19 pandemic highlighted the elderly as being especially vulnerable to increased disease severity and mortality, as well as an increased likelihood of long-term complications (2). In contrast, young individuals were often asymptomatic or had relatively mild-symptoms, and long-term complications were rare (2). While some hallmarks of aging are known to increase vulnerability to infection, for example immunosenescence, how the physiological state of “being old” determines the outcome of viral infections remains poorly understood.

The innate immune response is the first line of defense against invading viruses. In mammals, infected cells initiate a rapid innate immune response by detecting the DNA or RNA of the invading viruses with pathogen pattern recognition receptors (PRR), including the Toll-like receptors (TLR), retinoic acid-inducible gene I (RIG-I) like receptors (RLR), the cyclic GMP-AMP (cGAMP) synthase (cGAS), and nucleotide-binding oligomerization domain (NOD)-like receptors (NLR) (3). PRRs trigger the production of immune effectors, which include cytokines and chemokines (4). Among these immune effectors, the Type I interferon response plays a crucial role in regulating antiviral innate immune response. One important pathway that triggers the production of type I interferons (IFN-I) by RLRs includes RIG-I and melanoma differentiation-associated gene 5 (MDA5). Once activated, RIG-1 and MDA5 translocate to the outer membrane of mitochondria and bind to mitochondrial antiviral-signaling proteins (MAVS) to initiate a signaling cascade that facilitates cell death and removal of the infected cells (5).

*Caenorhabditis elegans* infection by Orsay virus (OV), a natural pathogen, is a novel and powerful genetic system to elucidate how aging impacts host-viral interactions, physiological response, and disease outcome. Cell culture lacks systemic-level responses seen *in vivo* and can lead to *in vitro* artifacts (6). Mechanisms preserving cell function at the molecular/subcellular level are difficult to discover *in vivo* within higher metazoans, as they are masked by cell death/replacement and immune clearance mechanisms that maintain tissue homeostasis; these issues are further confounded by compensatory and redundant mechanisms, which could mask the genetic contribution of underlying deficiencies. In contrast, the absence of an acquired immune system combined with an innate immune system that does not rely upon cell death or dedicated immune cells allows investigation into how non-immune cells maintain homeostasis *in vivo*. Although *C. elegans* has provided key insight into biological principles of aging (7, 8), how aging alters antiviral defense remains unexplored.

OV is a small, enteric single stranded RNA+ virus of the nodavirus family and the only known natural viral pathogen of *C. elegans* (9). Infection activates DRH-1, the ortholog of the DEAD/H-box Helicase and cytosolic pattern recognition receptor RIG-I to limit viral infection and pathogenesis (10-16). The OV genome is segmented into two parts: RNA1 encodes an RNA-dependent RNA polymerase (RdRP), and RNA2 encodes capsid and delta proteins (17-19). The *C. elegans* intestine consists of 20 non-renewable cells, which last the lifespan of the animal. OV typically only infects between two to six intestinal cells; loss of RNAi has been reported to increase the amount of viral protein within a cell without increasing the number of infected cells (20), which suggests RNAi acts solely within the infected cell to reduce levels of virus. In a separate pathway that is independent of RNAi, DRH-1 also activates ZIP-1 (a bHLH transcription factor) to induce the Intracellular Pathogen Response (IPR) (21, 22). The IPR constitutes a shared adaptive transcriptional response of ∼80 genes induced by OV, fungal pathogens, and some non-pathogenic forms of stress, including proteotoxicity following acute heat shock or inhibition of the proteosome (22, 23); the IPR has similarities to the Type I IFN response (23).

In the current study, we sought to gain mechanistic insight into how DRH-1 is activated by viral infection and how the antiviral activity of DRH-1 changes during aging. We found that SUMOylation of DRH-1 negatively regulated the IPR and facilitated viral infection. Sumoylation is a rapid and reversible post-translational modification that plays conflicting roles in viral replication and antiviral defenses in mammals (24). We found increased viral transcript levels, bloating within the intestinal lumen, and impaired induction of the IPR in animals lacking the SUMO isopeptidase *ulp-4* (orthologous to mammalian SENP7), which suggested a more severe infection. In the absence of *ulp-4*, basal levels of DRH-1 are reduced via increased degradation through the proteosome, consistent with findings that SUMOylated proteins can be degraded by the UPS (25). After infection, ULP-4 deSUMOylated DRH-1 to facilitate translocation to the mitochondrial outer membrane to activate the IPR; mutating DRH-1 lysine residues that are targets for SUMOylation rescued the impaired IPR and levels of DRH-1 protein. During aging, *ulp-*4 mRNA decreased and inducibility of the IPR was lost. Overexpression of a non-SUMOylatable isoform of DRH-1, specifically restored the inducibility of the IPR in older animals, which suggests that age-associated dysregulation of DRH-1 SUMOylation contributes to an increase in pathogenesis in older animals.

## Results

### The SUMO isopeptidase ULP-4 is required for induction of the intracellular pathogen response by viral infection

SUMOylation is a post-translational modification process resembling ubiquitination; the *C. elegans* genome encodes a single SUMO moiety (SMO-1), two E1 ligases (UBA-2, AOS-1), one E2 ligase (UBC-9) and one E3 enzyme (GEI-17), which conjugate SMO-1 onto proteins. Conversely, there are four SUMO isopeptidases that deconjugate SUMO from a target protein: ULP-1, ULP-2, ULP-4 and ULP-5. To begin to explore the roles of SUMOylation in the regulation of antiviral innate immunity, we inactivated components of the SUMOylation machinery by feeding RNAi and assessed the impact on the induction of the IPR after viral infection. Induction of *pals-5* expression (*pals-5::GFP*) is a canonical readout for induction of the IPR (15, 21, 26). Of the four SUMO isopeptidases, only loss of *ulp-4* significantly limited *pals-5p::GFP* induction after viral infection (Fig. 1*A*, *SI Appendix*, Fig. 1*A*), which was approximately 4-fold lower than induction in virally infected animals treated with empty vector RNAi (Fig. 1*B*). Conversely, inactivation of components that promote SUMO conjugation (*aos-1(RNAi), gei-17(RNAi), ubc-9(RNAi)*), or loss of SUMO itself (*smo-1(RNAi)*), significantly induced *pals-5p::GFP* after infection (Fig. 1*B*, *SI Appendix*, Fig. 1*A*). Inactivation of any component of the SUMO core machinery was not sufficient to induce *pals-5* in the absence of viral infection (Fig 1*A, SI Appendix*, Fig. 1*A*), suggesting that altered SUMOylation status plays a regulatory role in the response to viral infection. To confirm that loss of SUMO conjugation enhanced induction of the IPR, we tested whether *gei-17(tm2723)* deletion mutants recapitulated *gei-17(RNAi)*; viral infection of *gei-17* mutant animals induced *pals-5p::GFP* approximately 3-fold higher than induction in wildtype animals (*SI Appendix*, Fig. 1*B* and *C*). Next, we tested whether *ulp-4(tm3688)* deletion mutants recapitulated *ulp-4(RNAi)*; the relative induction of *pals-5* mRNA was approximately 5-fold less in the absence of *ulp-4* (Fig. 1*C*). Viral load was higher in the absence *of ulp-*4, either via RNAi or in null-mutant animals (Fig. 1*D,E*). Next we tested whether non-viral triggers that induce the IPR also required *ulp-4* (i.e., proteasome blockade or heat shock (15); *pals-5p::GFP*). Significantly, we found *ulp-4* is not required for the induction of *pals-5* by these two non-viral triggers (*SI Appendix*, Fig. 2*A-D*). We conclude that ULP-4 has an essential and specific role in activating an antiviral defense response after OV infection.

**Figure 1.**
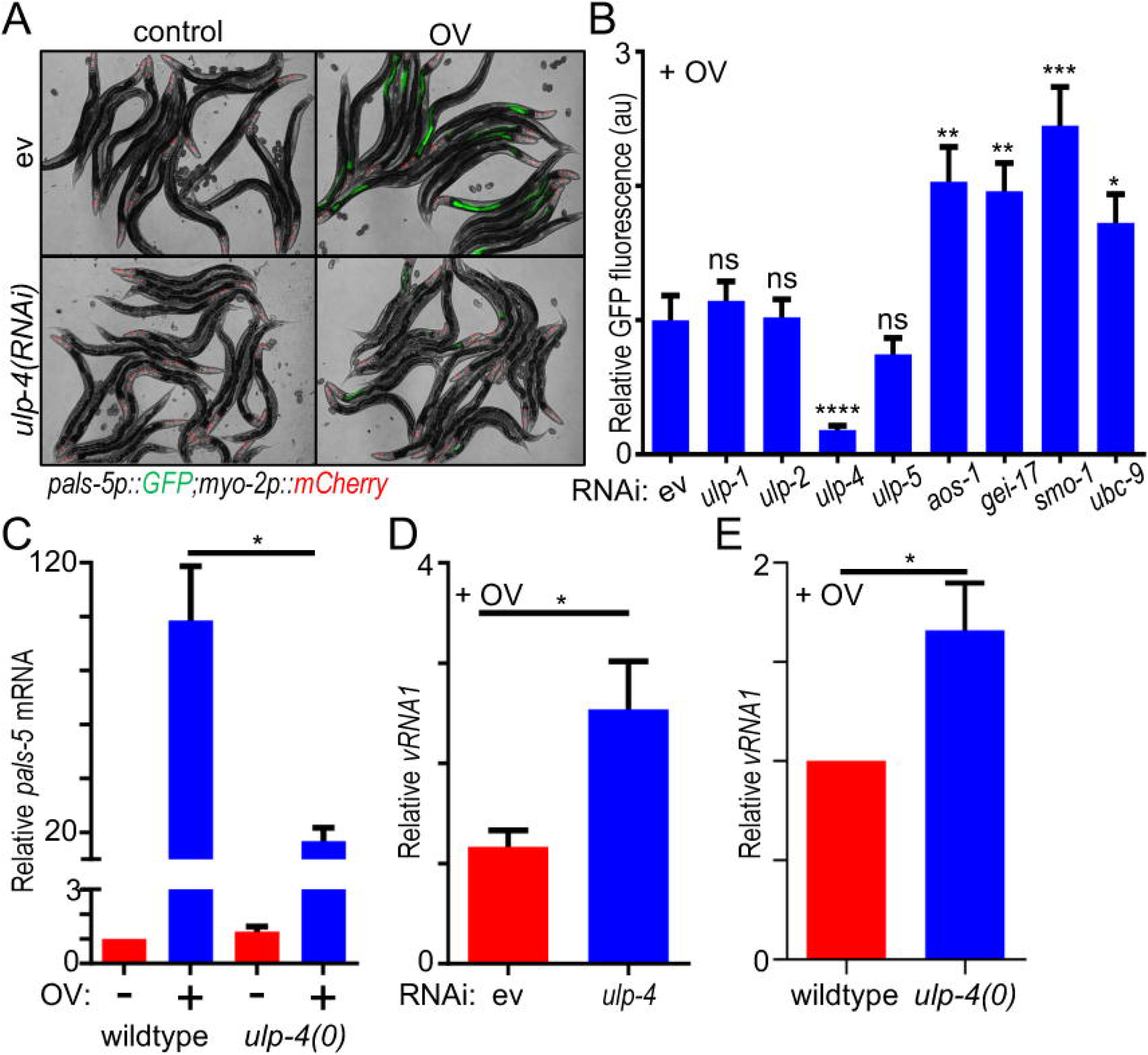
ULP-4 is required for induction of the intracellular pathogen response after viral infection. (*A*) Representative images of *pals-5p::GFP* expression after empty vector (ev) or *ulp-4* RNAi +/-viral infection. *myo-2p::mCherry* is a pharyngeal expressed co-injection marker to identify the *jyIs8[pals-5p::gfp]* transgene. Scale bar = 200 µm. (*B*) Quantification of *pals-5p::GFP* fluorescence intensity after inactivation of SUMOylation machinery + viral infection. Representative images +/-viral infection are in *SI Appendix* Fig. 1*A*. (*C*) RT-qPCR analysis of endogenous *pals-5* mRNA in either wildtype or *ulp-4(tm3688)* null mutant animals infected with virus. (*D*) RT-qPCR analysis of Orsay virus *RNA1* levels in (*A*). (*E*) RT-qPCR analysis of Orsay virus *RNA1* levels in (*C*). In all cases, the relative mean of 60 animals across three independent trials is shown. Error bars are the SEM. A two-tailed t test was used to calculate P-values; *P < 0.05, **P < 0.01, ***P < 0.001, ****P < 0.0001, ns = no significant difference.

**Figure 2.**
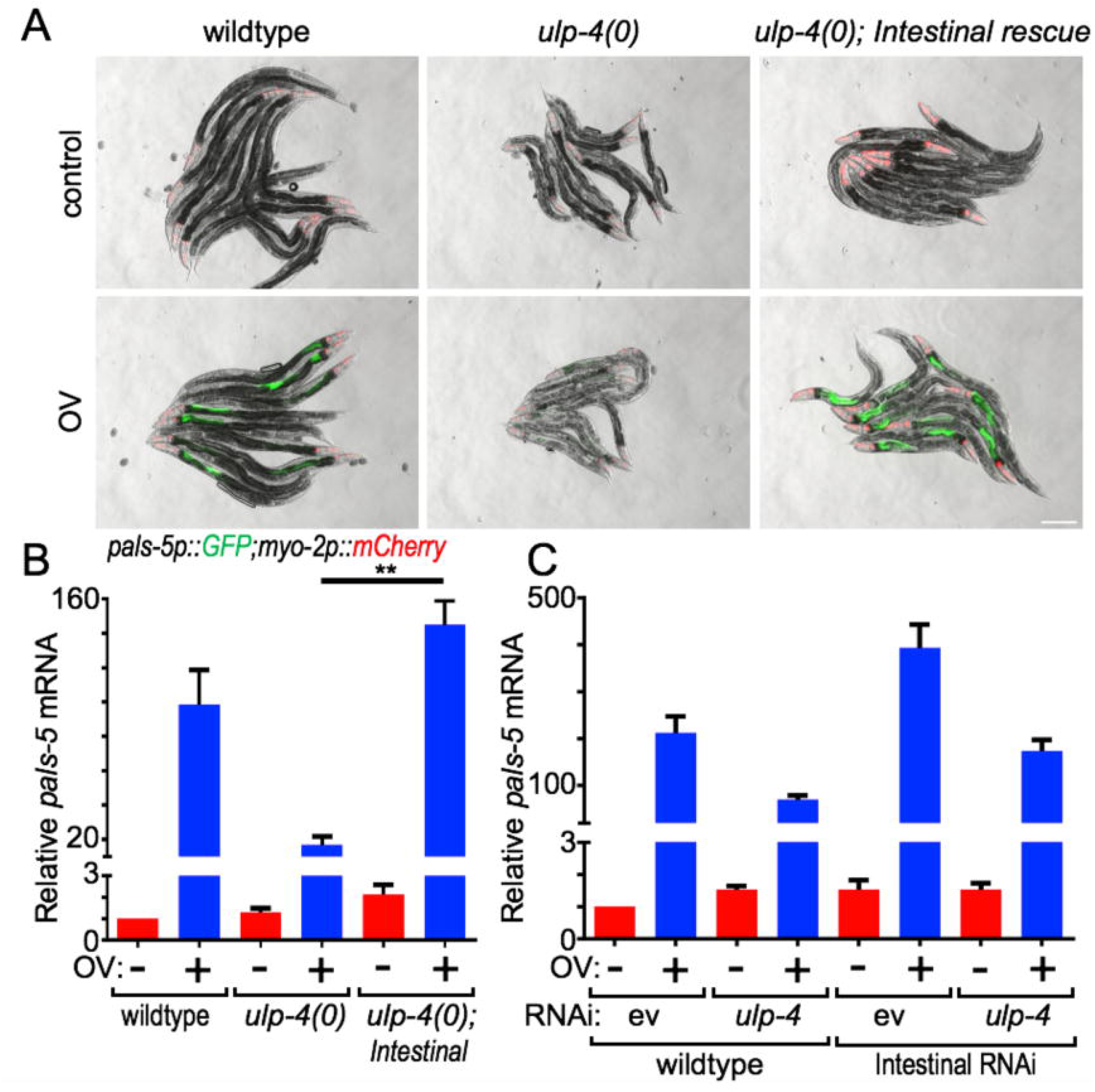
ULP-4 acts in the intestine to regulate the IPR upon viral infection. (*A*) Representative images of *pals-5p::GFP* expression +/-viral infection of either wildtype, *ulp-4(tm3688)* null, or *ulp-4(tm3688);artEx95(ges-1p::ULP-4)* animals. Scale bar = 200 µm. (*B*) RT-qPCR analysis of endogenous *pals-5* mRNA of samples in (*A*). (*C*) RT-qPCR analysis of endogenous *pals-5* mRNA +/-viral infection after systemic or intestine-specific inactivation of *ulp-4*. In all cases, values are the mean across three independent trials; error bars are SEM. A two-tailed t test was used to calculate P-values; *P < 0.05, **P < 0.01.

### ULP-4 acts in the intestine to regulate antiviral immune response

To explore whether ULP-4 regulates antiviral immune signaling cell autonomously within intestine or cell non-autonomously, we used tissue specific promoters to rescue *ulp-4* expression with the intestine (*ges-1p*), muscle (*myo-3p*), epidermis (*dpy-*7*p*) or nervous system (*rab-3p*) in the *ulp-4(tm3688)* null mutants, and then assessed whether *pals-5* induction was restored following viral infection. We found that only intestinal *ulp-4* expression (*ges-1p::ulp-4*) restored *pals-5* induction after viral infection, as measured by GFP fluorescence *(pals-5p::GFP*) and levels of endogenous mRNA (Fig. 2*A,B*; *SI Appendix*, Fig. 3*A,B*). Next, we conducted the converse experiment using intestinal specific RNAi. Intestinal loss of *ulp-4* impaired the inducibility of the IPR to a comparable extent as systemic inactivation (Fig. 2*C*). We conclude that ULP-4 functions within the intestine to regulate induction of the IPR in response to viral infection, which is consistent with either a cell autonomous or localized intestinal response to an enteric viral infection.

**Figure 3.**
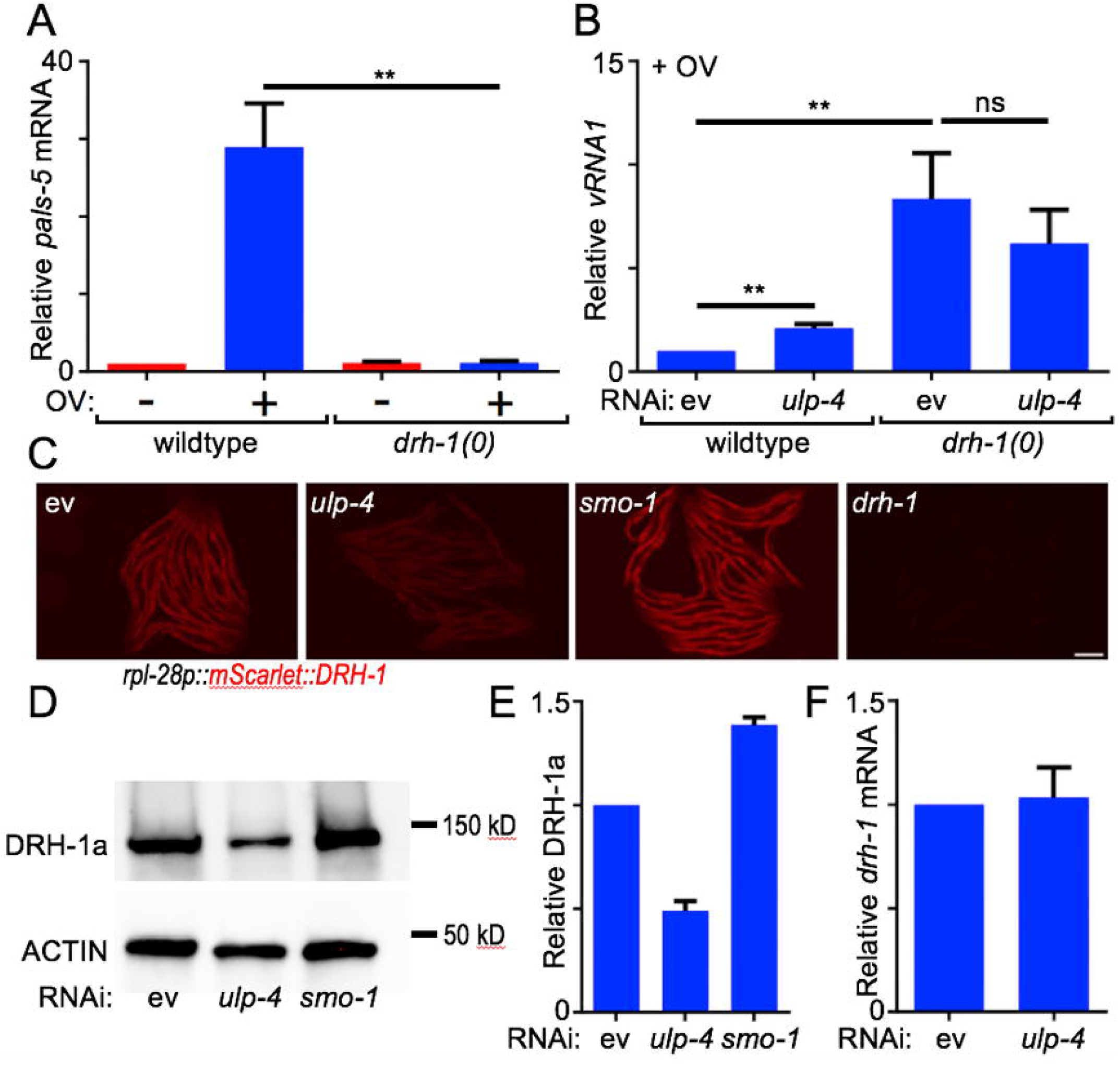
ULP-4 promotes the protein stability of DRH-1 through deSUMOylation. (*A*) RT-qPCR analysis of endogenous *pals-5* in wildtype and *drh-1(ok3495)* null mutants +/-viral infection. (*B*) RT-qPCR analysis of virus *RNA1* levels in wildtype and *drh-1(ok3495)* null animals treated with either empty vector or *ulp-4(RNAi)* followed by viral infection. (*C*) Representative images of *mScarlet::DRH-1* expression in uninfected animals after RNAi to empty vector, *ulp-4, smo-1*, or *drh-1*, respectively. Scale bar = 200 µm. (*D*) Representative immunoblot of conditions in panel (*C*). (*E*) Quantification of DRH-1 protein levels. (*F*) RT-qPCR analysis of endogenous *drh-1* after empty vector or *ulp-4(RNAi)*. In all cases, values are the mean across three independent trials; error bars are SEM. A two-tailed t test was used to calculate P-values; *P < 0.05, **P < 0.01,***P < 0.001.

### ULP-4 acts in a common pathway with DRH-1 to limit viral infection

To begin to understand how ULP-4 activity regulates the IPR in response to viral infection in young animals, we first determined whether ULP-4 and DRH-1 acted within a common genetic pathway. Consistent with previous reports that DRH-1 signaling is necessary for the induction of the IPR (15), we found that *drh-1(ok3495)* null mutant animals failed to induce the IPR following viral infection (*pals-5p::GFP*; Fig. 3*A*). Next, we found that simultaneous loss of both *ulp-4* and *drh-1* did not produce higher viral loads in infected animals: inactivating *ulp-4* in otherwise wildtype animals resulted in a small but significant increase in viral loads, as measured by levels of Orsay virus RNA1 (Fig. 3*B*). In contrast, infected *drh-1(ok3495)* null mutant animals had viral loads ∼8.4-fold higher than wildtype; there was no significant additivity when both *ulp-4* and *drh-1* were lost (Fig. 3*B*). It is unlikely that a lack of additivity was due to a threshold effect, as impairing other antiviral innate defenses, such as RNAi, can produce viral loads several orders of magnitude than those observed in wildtype (9). The relative increase in viral loads were negatively correlated with the level of IPR induction after loss of either *ulp-4* or *drh-1* (compare Fig. 2*C* & 3*B* and Fig. 3*A* & 3*B* for loss of *ulp-4* and *drh-1*, respectively). This suggests that while ULP-4 and DRH-1 have common antiviral functions, in the absence of *ulp-4*, DRH-1 retains some activity.

### ULP-4 prevents proteosomal degradation of DRH-1

We tested whether modulation of SUMOylation *in vivo* would alter the stability of DRH-1 protein. In the absence of virus, loss of *ulp-4* was sufficient to significantly diminish DRH-1 protein levels throughout the animal; concordantly, loss of *smo-1* resulted in higher levels of DRH-1 (*mScarlet::DRH-1*; Fig. 3*C*). Red fluorescence was specific to the *mScarlet::DRH-1* transgene, as *drh-1(RNAi)* reduced fluorescence to almost undetectable levels (Fig. 3*C*). To confirm this finding, we conducted quantitative immunoblotting of DRH-1 after inactivation of either *ulp-4* or *smo-1*; basal levels of DRH-1 significantly decreased after *ulp-4(RNAi)* and increased after *smo-1(RNAi)* treatment (Fig. 3*D* and *E*). As expected, ULP-4 regulation of DRH-1 was strictly post-transcriptional, as level of *drh-1* mRNA were unchanged after inactivation of *ulp-4* (Fig. 3*F*). To gain insight into how ULP-4 regulated levels of DRH-1, we inactivated either the proteosome (*ubq-2(RNAi)*) or macroautophagy (*lgg-1(RNAi)*, hereafter referred to as autophagy), in conjunction with loss of *ulp-4* or *smo-1*, and assessed levels of DRH-1. *ubq-2* and *lgg-1* encode the orthologs of UBA52/ubiquitin and Atg8/LC3, which are required for proteosome function and macroautophagy, respectively (27, 28). Loss of *ubq-2* was sufficient to significantly increase fluorescence of the *mScarlet::DRH-1* transgene and DRH-1 protein levels in animals lacking *ulp-4, smo-1*, or that were otherwise wildtype. In contrast, loss of autophagy did not significantly alter levels of DRH-1 (*SI Appendix*, Fig. 4*A-C*). Next, we tested whether infection was sufficient to stabilize DRH-1. We found that DRH-1 levels were unchanged by viral infection (*SI Appendix*, Fig. 5*A,B*); however, OV typically only infects a few intestinal cells (20), and OV may only stabilize DRH-1 within infected cells (assessed below). Collectively, we conclude that ULP-4 acts upstream to stabilize DRH-1 by limiting SUMOylation, either directly or indirectly, and that this stability is required for the subsequent induction of IPR in response to viral infection.

**Figure 4.**
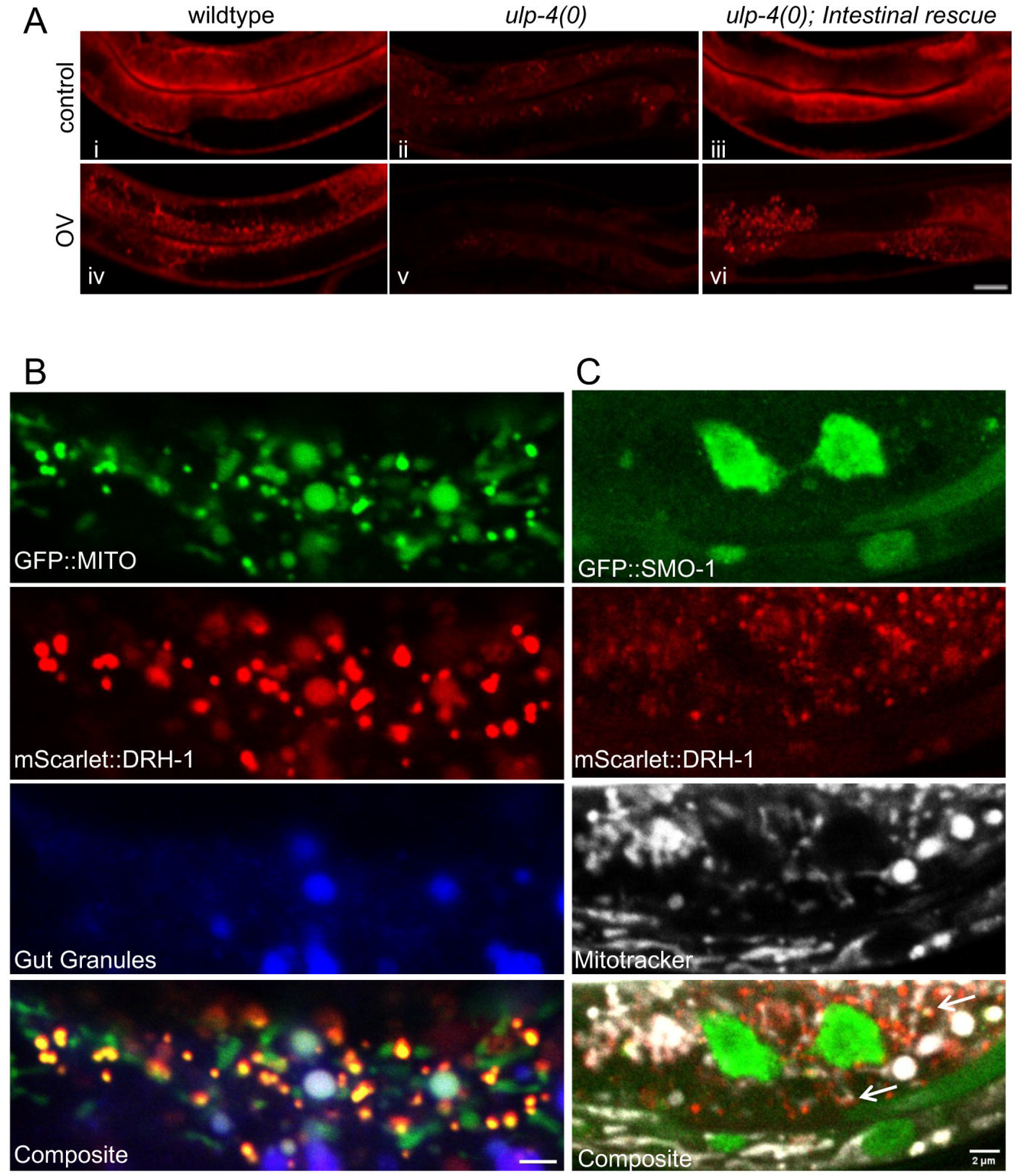
NonSUMOylated DRH-1 translocates to the mitochondria upon viral infection. (*A*) Representative images of mScarlet::DRH-1 in the intestinal region of wildtype, *ulp-4(tm3688)*, or *ulp-4(tm3688);artEx95(ges-1p::ULP-4)* of uninfected (*i-iii*) or virally infected animals (*iv-vi*). Scale bar = 20 µm. (*B*) Representative images of an intestinal cell of a *mScarlet::DRH-1;mito-GFP* animal infected with virus; lysosomal related organelles/gut granules are indicated (405-nm blue channel). Scale bar = 2 µm. (*C*) Representative image of an intestinal cell of a *mScarlet::DRH-1;SMO-1::GFP* animal infected with virus; mitochondria are indicated (Mitotracker, Far Red). Scale bar = 2 µm.

**Figure 5.**
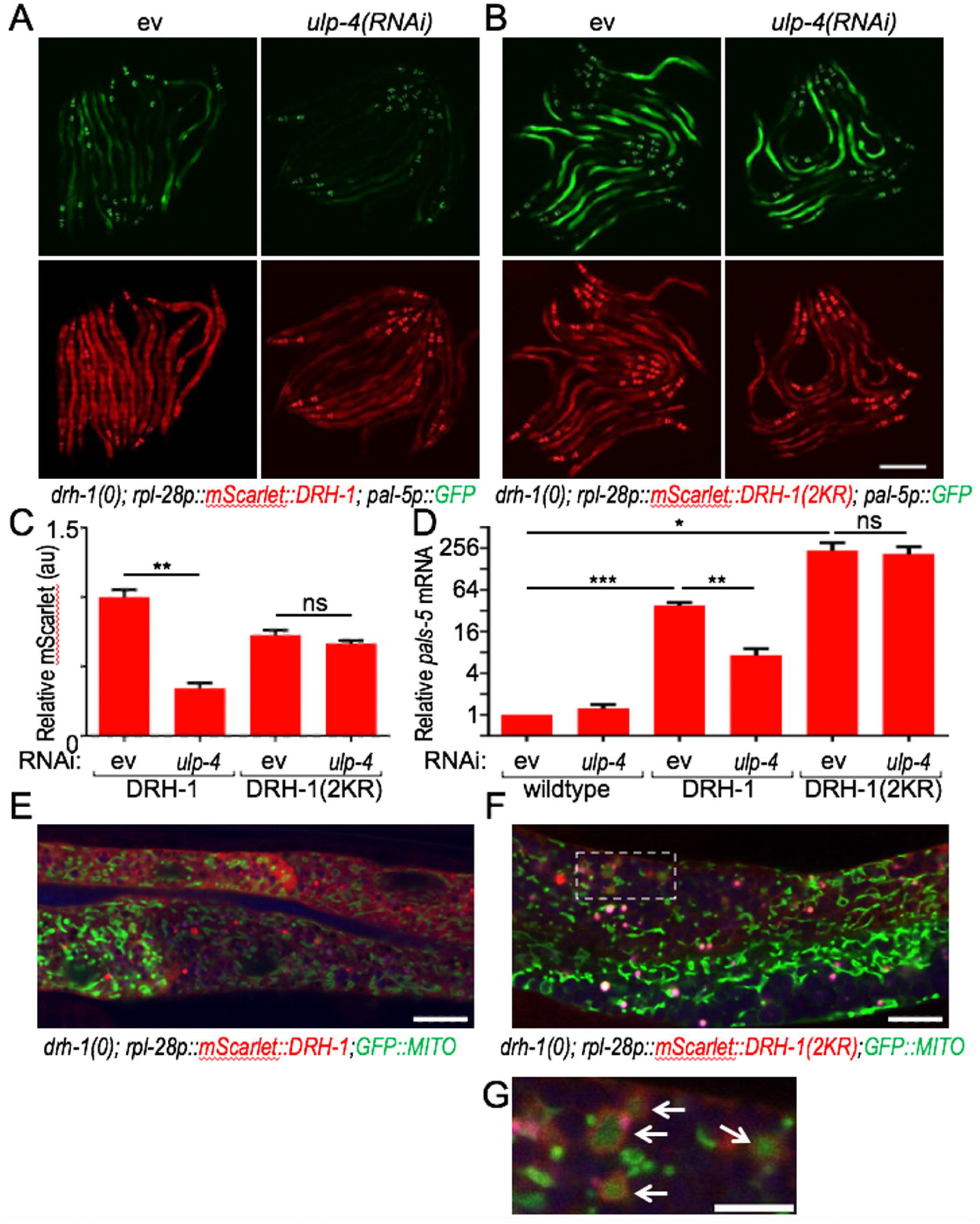
deSUMOylation of DRH-1 activates antiviral defense. (*A,B*) Representative images of *pals-5p::GFP* and *mScarlet* expression in *drh-1(ok3495)* plus somatic overexpression of either wildtype DRH-1 (*rpl-28p::mScarlet::DRH-1(wt), A*) or nonSUMOylatable DRH-1 (*rpl-28p::mScarlet::DRH-1(2KR), B*) treated with either empty vector or *ulp-4(RNAi)*. Scale bar = 200 µm. (*C*) Quantification of mScarlet::DRH-1 in (*A,B*). (*D*) RT-qPCR analysis of endogenous *pals-5* of samples from (*A,B*) and *JyIs8* wildtype control. (*E-G*) Representative composite images of uninfected L4 animals expressing *mScarlet::DRH-1(WT)* (*E*); *GFP::mito* or *mScarlet::DRH-1(2KR); GFP::mito* (*F,G*) respectively. Scale bar = 2 µm. Inset scale bar = 5 µm. For individual panels see *SI Appendix*, Fig. 7. In all cases, values are the relative mean from three independent trials; error bars are the SEM. A two-tailed t test was used to calculate P-values; *P < 0.05, **P < 0.01,***P < 0.001, ns = no significant difference.

### NonSUMOylated DRH-1 translocates to mitochondria upon viral infection

To further elucidate how SUMOylation regulates DRH-1, we assessed localization of DRH-1 following viral infection. Consistent with the findings of others (16), viral infection initiated a shift in DRH-1 from a diffuse cytosolic to the formation of puncta (Fig. 4*A*). In the absence of *ulp-4*, DRH-1 levels (mScarlet::DRH-1) were almost undetectable, but restoring *ulp-4* expression within the intestine was sufficient to restore DRH-1 induction and the puncta formation following viral infection (Fig. 4*A*), which suggests ULP-4 acts cell autonomously, perhaps directly upon DRH-1, to regulate the IPR. To determine the sub-cellular localization of DRH-1 containing puncta, we co-expressed *mScarlet::DRH-1* with GFP fused to a mitochondrial localization tag (*ges-1p::GFP(mit)*). Following viral infection, we observed that DRH-1 primarily co-localized with mitochondria within infected intestinal cells (Fig. 4*B*), To a much lesser extent, some overlap with auto-fluorescent gut granules were also observed (Fig. 4*B*), which have previously been shown to be lysosomal related organelles (LROs) (29). Next, we determined whether the SUMOylation status of DRH-1 dictated subcellular localization. Using transgenic animals co-expressing *GFP::SMO-1, mScarlet::GFP* and treated with Mitotracker, we found little to no overlap between the SUMO moiety (SMO-1) and DRH-1 at puncta that form during viral infection (Fig. 4*C*). Collectively, our findings indicate that unSUMOylated DRH-1 translocates to mitochondria following viral infection, and this is mediated by the deSUMOylating activity of ULP-4.

### deSUMOylation of DRH-1 activates antiviral defense

We posit ULP-4 may directly deSUMOylate DRH-1 to activate the IPR. SUMOylation frequently occurs at a consensus sequence (ψKxE) We used four different prediction tools to identify potential SUMOylation sites in DRH-1 (described in methods and (30-33)). Among the 60 lysine residues in DRH-1, all four algorithms only predicted K647 and K731 as targets for SUMOylation with high confidence (*SI Appendix*, Fig. 6*A*; *Dataset S1*). While K647 does not fall within a specific domain, K731 is located within the helicase domain. All RIG-I like receptors (RLR) have a central helicase domain and a carboxy-terminal domain (CTD), which work together to detect immunostimulatory RNAs (34). To functionally test whether K647/731 are the SUMOylation sites of DRH-1, both were mutated to arginine, which prevents SUMOylation at those residues. Transgenic animals were generated that express either the mutated (2KR) or wildtype isoform of DRH-1a throughout the soma (*rpl-28p::mScarlet::DRH-1(2KR)*); transgenes were expressed in *drh-1(ok3495)* null mutants at similar levels, approximately 4-fold higher than endogenous *drh-1* (*SI Appendix*, Fig. 6*B*). Protein levels of DRH-1 and DRH-1(2KR) were also similar *in vivo* (Fig. 5*A,B,C*). Consistent with the findings of others (16), overexpression of wildtype *drh-1* was sufficient to modestly activate the IPR *in vivo* in the absence of virus (*pals-5p::GFP*; Fig. 5*A*). In contrast, DRH-1(2KR) animals robustly induced *pals-5p::GFP* fluorescence (Fig. 5*B*), despite having slightly lower levels of red fluorescence (mScarlet::DRH-1(2KR); Fig. 5*C*). Loss of *ulp-4* suppressed the induction of the IPR in animals overexpressing the wildtype isoform of DRH-1 (Fig. 5*A*). In contrast, loss of *ulp-4* had no effect on the induction in animals expressing *drh-1(2KR)* (Fig. 5*B*). Similar results were found with expression of endogenous *pals-5* mRNA (Fig. 5*D*), and of another gene within the IPR (*sdz-6*; *SI Appendix*, Fig. 6*C*). Lastly, we identified the subcellular distribution of DRH-1 and DRH-1(2KR) under basal conditions. As expected, in the absence of viral infection, wildtype DRH-1 was predominantly diffuse throughout the cytosol but overexpression led to the formation of some puncta (Fig. 5*E*, SI *Appendix*, Fig. 7), which is consistent with modest induction of the IPR. In contrast, mScarlet::DRH-1(2KR) was predominantly localized in puncta either surrounding mitochondria or at LROs neighboring mitochondria (Fig. *5F,G*, *SI Appendix*, Fig. 7); the latter observation suggests DRH-1 activation may have antiviral activity through the coupling mitochondrial and lysosomal activity. Collectively, we conclude that SUMOylation normally limits activation of DRH-1 and that upon viral infection ULP-4 deSUMOylates DRH-1 allowing translocation to the mitochondria to activate innate antiviral defenses.

**Figure 6.**
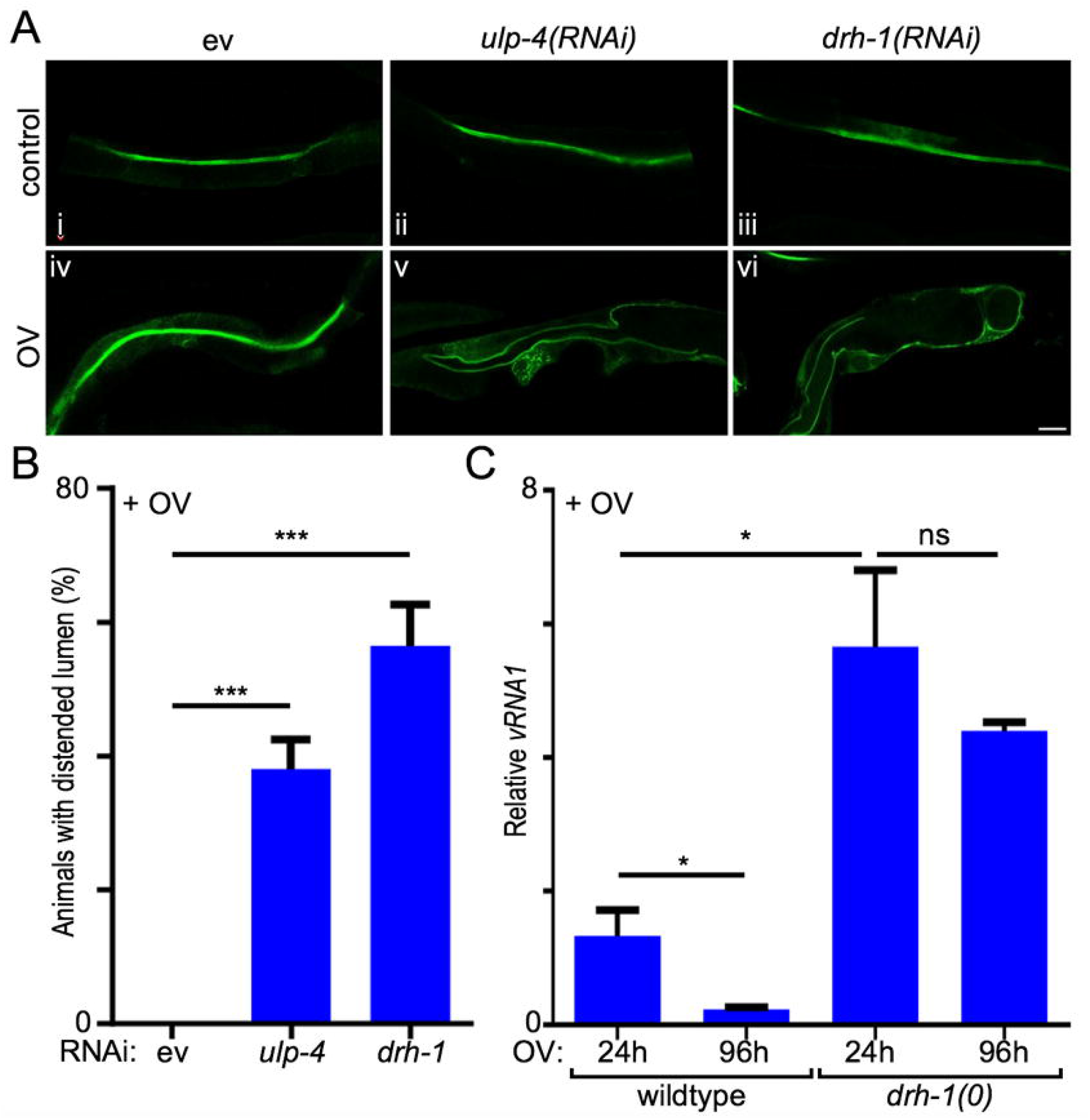
Antiviral defense through ULP-4 and DRH-1 limit viral pathogenesis. (*A*) Representative images of animals expressing the *ACT-5::GFP* apical membrane marker treated with either empty vector, *ulp-4(RNAi)*, or *drh-1(RNAi)*, respectively, +/-viral infection. Scale bar = 20 µm. (*B*) Quantification of the portion of a population with an enlarged lumen. (*C*) RT-qPCR quantification of Orsay virus *RNA1* in wildtype or *drh-1(ok3495)* null mutant animals infected with virus. In all cases, values are the mean of three independent trials; error bars are the SEM. A two-tailed t test was used to calculate P-values; ***P < 0.001, *P < 0.05, ns = no significant difference.

**Figure 7.**
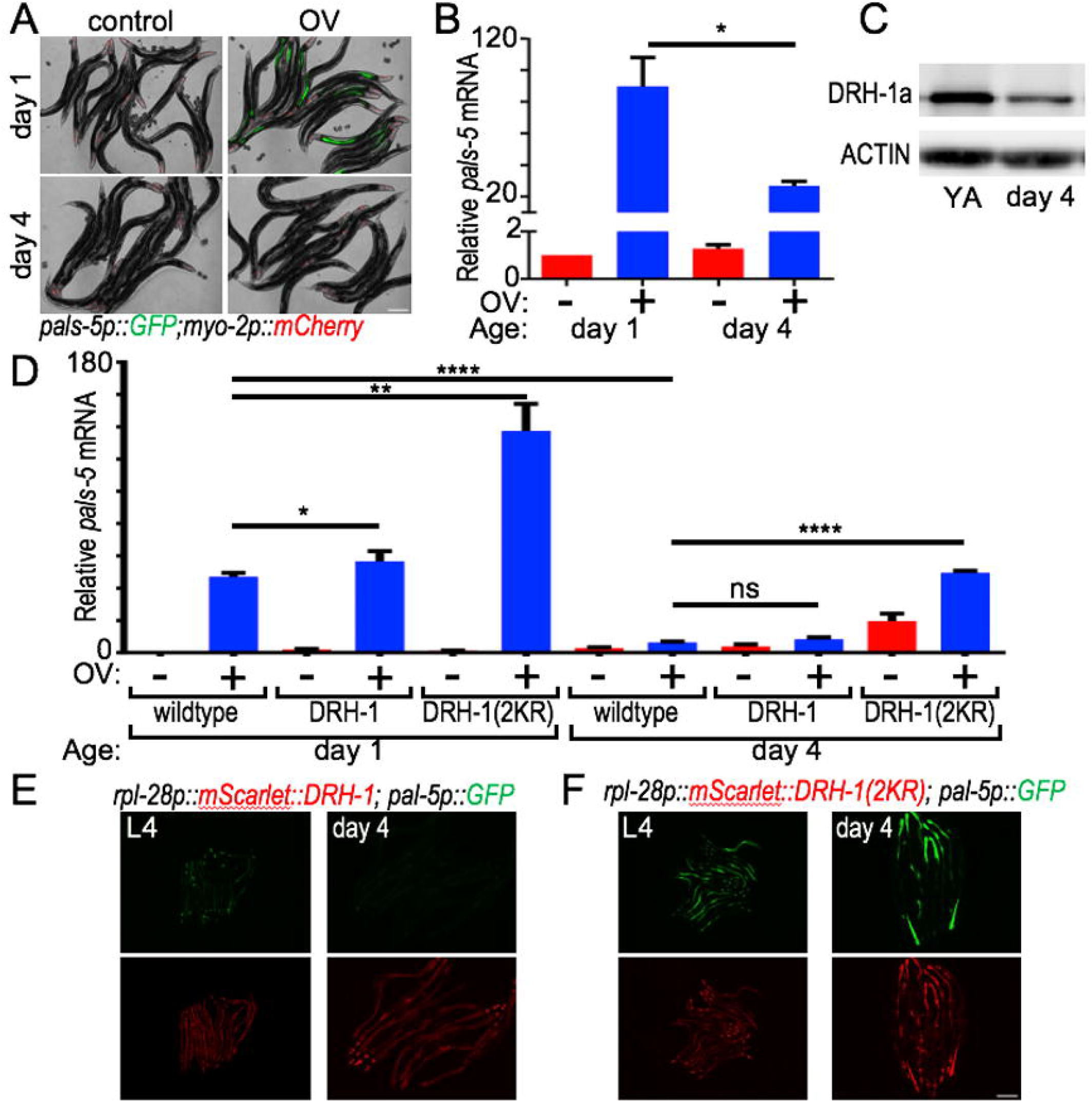
The IPR declines during aging due to dysregulation of DRH-1 SUMOylation. (*A*) Representative images of *pals-5p::GFP* expression +/-viral infection. Viral infection was initiated at L4 or day 3 of adulthood, and images were obtained 24 hours later. Scale bar = 200 µm. (*B*) Quantification of GFP fluorescence intensity of (*A*). (*C*) Representative immunoblot of mScarlet::DRH-1 levels in young adult (YA) or day 4 adult animals. (*D*) RT-qPCR analysis of endogenous *pals-5* +/-viral infection at different ages in wildtype, or *drh-1(ok3495)* null mutant animals with restored intestinal expression (*ges-1p*) of either wildtype DRH-1 (*mScarlet::DRH-1(wt)*) or nonSUMOylatable DRH-1 (*mScarlet::DRH-1(2KR)*). (*E,F*) Representative images of *pals-5p::GFP* expression at different ages in *drh-1(ok3495)* null mutant animals with restored somatic expression (*rpl-28p*) of either wildtype DRH-1 (*mScarlet::DRH-1(wt)*) or nonSUMOylatable DRH-1 (*mScarlet::DRH-1(2KR)*). In all cases, values are the relative mean of 60 animals across three independent trials; error bars are the SEM. A two-tailed t test was used to calculate P-values; *P < 0.05, **P < 0.01, **P < 0.0001, ns = no significant difference.

### Animals with compromised antiviral defense fail to clear virus and show signs of pathogenesis

To test the physiological consequences of *ulp-4* or *drh-1* loss, we assessed intestinal status using a tagged GFP that localizes at the apical membrane of the intestine to mark the lumen. In the absence of viral infection loss of either *ulp-4* or *drh-1* had no effect on the lumen (Fig. 6*Ai-iii*). Similarly, viral infection was not sufficient to cause obvious changes to the intestinal lumen (Fig. 6*Aiv*). However, in the absence of either *ulp-4* or *drh-1* viral infection produced signs of intestinal bloating in a significant portion of infected animals (Fig. 6*Av,vi,B*). Intestinal bloating has previously been shown to occur during infection with bacterial pathogens and is indicative of increased pathogenesis (35). Next, we tested whether loss of DRH-1 impacted viral load over time, a surrogate measure of viral clearance. Wildtype animals had a significant reduction of viral levels four days post-infection. In contrast, animals lacking *drh-1* not only have higher viral loads than wildtype, but *drh-1* null mutant animals also failed to significantly decrease viral loads over time (Fig. 6*C*). Thus, ULP-4 and DRH-1 play a critical role in activating the IPR in response to viral infection and that loss of this response results in increased pathogenesis and an inability to clear the virus.

### Antiviral response declines during aging due to dysregulation of DRH-1 SUMOylation

Aging results in the diminished inducibility of a myriad of adaptive transcriptional responses that maintain homeostasis following stress (36-38). We tested whether the inducibility of the IPR declined during normal aging and the role of SUMOylation. By the 4^th^ day of adulthood, the induction of the IPR by viral infection was severely impaired in wildtype animals (*pals-5p::GFP, pals-5* mRNA; Fig. 7*A,B*). At this age non-viral triggers of diverse forms of stress response are also known to be impaired (39-43) and global patterns of hyper-SUMOylation within the proteome have been reported (44). We found that basal levels of DRH-1 are decreased by the 4^th^ day of adulthood (Fig. 7*C*). We tested whether DRH-1 decreased due to degradation via the proteosome. Inactivation of the proteosome (*ubq-2(RNAi)*) increased levels of DRH-1 in older animals (*SI Appendix*, Fig. 8). However, overexpression of DRH-1 either within intestine or throughout the soma was not sufficient to restore inducibility of the IPR in older animals (Fig. 7*D,E*, respectively). This suggests that it is the proper regulation of DRH-1, but not levels of DRH-1 *per se*, that may breakdown during aging. We first tested this idea through inhibition of SUMOylation; we inactivated the sole SUMO moiety (*smo-1(RNAi)*) and tested whether this was sufficient to restore the inducibility of the IPR in older animals. In both young and old animals, inhibition of *smo-1* was sufficient to enhance the induction of the IPR after viral infection, but failed to fully restore the IPR (*SI Appendix*, Fig. 9*A,B*). While inhibition of *smo-1* elevated DRH-1 levels (*SI Appendix*, Fig. 9*C,D*), *smo-1(RNAi)* is likely to have off-target effects that could negatively impact overall cellular physiology and it is unclear how quickly inactivation of *smo-1* mRNA would diminish levels of conjugated and unconjugated SMO-1 in young or old animals. Therefore, we examined whether levels of *ulp-4* were altered in older animals and found that *ulp-4* mRNA significantly declined by day 4 of adulthood (*SI Appendix*, Fig. 9*E*), which suggests that the age-associated decline of ULP-4 compromised the IPR by the failure to remove SUMO from DRH-1. We tested this possibility by overexpressing *drh-1(2KR)* either throughout the intestine or soma. Unlike wildtype DRH-1, which failed to restore any inducibility of the IPR following viral infection, intestinal or somatic overexpression of *drh-1(2KR)* restored a level of IPR inducibility comparable to those seen in younger wildtype animals (Fig. 7*D,F*, respectively). We conclude that aged animals are unable to induce the IPR, at least in part, due to dysregulation of DRH-1 through increased SUMO conjugation.

**Figure 8.**
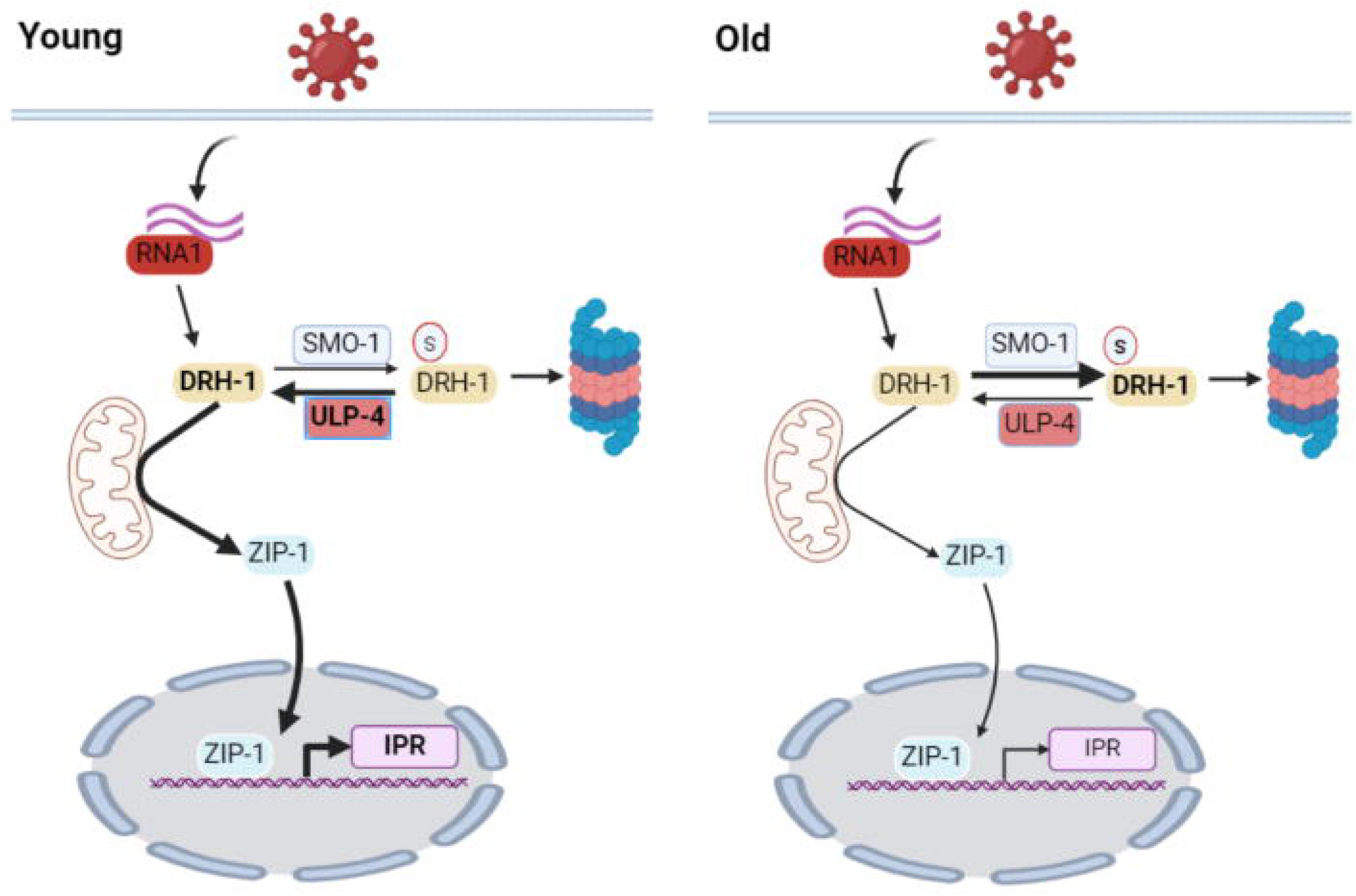
Model of dysregulation of SUMO conjugation to DRH-1 during aging, which compromises antiviral innate immunity to promote pathogenesis.

## Discussion

### ULP-4 deSUMOylates DRH-1 to prevent proteosomal degradation

In mammals, the deconjugating activity of SENP family members are specific: SENP1&2 remove all SUMO moieties (i.e., SUMO1-5) (45). In contrast, SENP6&7 are only responsible for removing SUMO2/3 (45-48). SUMO2/3 form covalent polymers that are recognized by SUMO-targeted ubiquitin ligases (STUbLs), which conjugate ubiquitin to form hybrid chains (49). SUMO-Ub hybrid chains target substrates for proteasomal degradation, but can also mediate nonproteolytic signaling (50-52). In contrast, SUMO1 cannot form monomeric chains (53). These structural differences are thought to be a major driver of the functional differences between SUMO1 and SUMO2/3 (49, 54). *C. elegans* SMO-1 has similarities to both SUMO1 and SUMO2/3: the sequence is more similar to SUMO1 but the electrostatic surface features are closer to SUMO2/3 (55). SMO-1 conjugation is thought to execute all of the functions of mammalian SUMO1 and SUMO2/3, including the formation of novel non-covalent protein-protein interactions through typical SUMO Interacting Motifs (SIMs) (55). SMO-1 conjugation has been shown to alter protein-protein interactions, initiate shifts in subcellular localization, and regulate chromatin and transcription factors (56-58). For instance, we previously found that SUMOylation of the heat shock transcription factor (HSF-1) limits the activation of the heat shock response in *C. elegans* (59).

Our work establishes that ULP-4, orthologous to SENP7, prevents the SMO-1 mediated proteosomal degradation of DRH-1, which acts as a regulatory mechanism that limits antiviral innate immunity (Fig. 8). In mammals, the role of SUMOylation in viral innate immunity through activation of the type I IFN response is complicated. RIG-1 has been shown to be SUMOylated by SUMO1 in response to RNA viral infection, which stabilizes RIG-1 to enhance IFN-I production by increasing interaction with <u>M</u>itochondrial <u>A</u>nti<u>v</u>iral-<u>S</u>ignaling (MAVS) proteins (60, 61). In stark contrast, SUMO2 and SUMO3 have previously been shown to act redundantly through an unknown mechanism to potently inhibit the activation of the Type I IFN response (62). Our results hint that the Type I IFN response may be negatively or positively regulated by SUMO2/3 or SUMO1, respectively, by converging upon RIG-I. Alternatively, the opposing roles of SUMO1 and SUMO2/3 in regulating antiviral innate immunity may have diverged; mammalian SENP isopeptidases also have opposing roles in viral innate immunity. SENP1 deconjugates SUMO1 to negatively regulate antiviral immunity by deSUMOylation of MAVS and activation of IRF3 (63). In contrast, SENP7 deSUMOylates cGAS, which induces expression of IRF3-responsive genes (64).

### Mitochondria are evolutionarily conserved antiviral signaling hubs

Mitochondria act as subcellular signaling hubs in response to stress, including viral infections (65). The type of response is dictated both by the nature and threshold of the stress; for example, low of levels of stress within the mitochondrial proteome activate the mitochondrial unfolded protein response (UPR^mito^), a molecular mechanism to restore homeostasis that preserves the cell. Increasing levels of stress activate subcellular responses including mitochondrial fusion to dilute damage and fission to remove damage that gets sequestered at the ends of filament networks, which can progress to mitophagy (65, 66). In mammals, chronic or high levels of stress induce programmed cell death pathways to preserve tissue homeostasis; the most well studied is apoptosis, which is triggered from mitochondria (67). In mammals, mitochondria also function as a platform for the innate immune response to infection by RNA viruses. Innate immunity in mammals relies on MAVS proteins on the outer mitochondrial membrane. Viral RNA binds to RIG-I and related RLRs to activate viral innate immune signaling (5, 34, 68, 69). Despite the lack of a MAVS ortholog in *C. elegans*, an analogous response is occurring, as DRH-1 translocates to the mitochondria after viral infection (Fig. 4) and mitochondria still serve as the focal point for initiating antiviral response. Thus, mitochondria play a central role in antiviral innate immunity, even though evolutionary pressure between viruses and hosts may have favored a divergence in the specific regulation of antiviral components.

A role linking SENP7 and mitochondrial function in mammals has not been described, but emerging results in *C. elegans* support a possible link. The handful of *C. elegans* studies on ULP-4 have implicated a key role for coordinating stress response initiated in mitochondria (70, 71). First, ULP-4 has been shown to regulate the first enzyme of the mevalonate pathway, which undergoes age-dependent SUMOylation and correlated with a shift in ULP-4 subcellular localization from the cytosol to the mitochondria in non-intestinal tissues (71). In another study, ULP-4 was shown to be a positive regulator of DVE-1 and ATFS-1, two transcription factors that are the primary effectors of the UPR^mito^ (70). In the absence of *ulp-4*, DVE-1 failed to translocate to the nucleus and the induction of the UPR^mito^ is impaired (70). Viral targeting of mitochondria generates stress that induces retrograde signals (72, 73), and our findings that ULP-4 deSUMOylates DRH-1 to facilitate mitochondrial localization and activation of the IPR suggest this may be conserved in *C. elegans*. We find it intriguing that ULP-4 regulates both the UPR^mito^ and IPR, two mitochondrial signal transduction pathways that maintain homeostasis. How target selectivity of SENPs is obtained remains a mystery, but given the relatively small number of SUMO isopeptidase genes it has been postulated that groups of proteins functionally and physically linked are regulated by the same SENP (74). For example, SUMO often targets multiple proteins within a complex, or within a pathway (75, 76). It will be important in future studies to identify how ULP/SENP target selectivity is obtained to maintain homeostasis in animals experiencing diverse forms of stress linked to mitochondria.

### Dysregulated SUMO conjugation of DRH-1 during aging compromises antiviral innate immunity to promote pathogenesis

*C. elegans* provides an opportunity to determine how antiviral responses in non-immune cells change with age, which would be difficult to discover *in vivo* in more complex metazoans. While incredibly complex, immunosenescence in mammals is well-described and can be most simply defined as the gradual deterioration of both the acquired and innate immune systems during aging. Immunosenescence is also associated with the onset of a chronic, systemic inflammatory state (i.e., inflammaging) (6, 77-79). The study of chronic viral infections in the elderly that are attributed to declining innate immunity has largely focused on neutrophils, monocytes, macrophages, natural killer cells, natural killer T (NKT) cells and dendritic cells (80-83). Our findings that loss of the IPR results in a chronic viral infection and produces signs of pathogenesis in *C. elegans* is consistent with the possibility that the aging of non-immune cells in higher vertebrates is an additional level of immunosenescence.

Our discovery that the inducibility of the innate immune response declines due to the loss of ULP-4-mediated deSUMOylation of DRH-1 represents a novel age-associated breakdown in antiviral defense. In *C. elegans*, several published studies -including two from our laboratory, have implicated either SMO-1 or orthologs of SENPs in aging or the regulation of pathways associated with healthy aging (44, 59, 70, 71, 84), but viral innate immunity had not been previously investigated. Emerging evidence suggests that an age-associated breakdown in proper SUMOylation of the proteome contributes to aging and age-associated disease in humans. SUMOylation plays a key role in protein quality control and the maintenance of proteostasis and has been implicated in nearly all age-associated neurological disorders (49). Our discoveries suggest a possible link between the age-associated loss of antiviral defense and a shift in the physiological state of non-immune cells, which may contribute to the accumulation of endogenous retroviral elements and increased viral pathogenesis in the elderly.

## Materials and Methods

*C. elegans* strains (*SI Appendix*, Table S1) were maintained on nematode growth medium (NGM) plates seeded with OP50 Escherichia coli. Worms were maintained at 20°C unless otherwise noted. All QPCR primers used in this study are listed in *SI Appendix*, Tables S2. RNAi and tissue-specific RNAi were performed via the feeding method. All infection assays were performed on developmentally synchronized animals. Orsay virus infections were performed using virus from the same batch of virus filtrate. All Constructs were generated with the NEBuilder HiFi DNA Assembly Cloning Kit (NEB #E5520). Quantification of *pals-5p::GFP* and *mScarlet::DRH-1* reporter fluorescence was performed in Fiji. All subcellular imaging was captured with a Leica DMi8 confocal microscope. The rabbit polyclonal anti-RFP antibody (Rockland # 600-401-379) was used for DRH-1 immunoblotting. All statistical analysis was performed with GraphPad Prism. A two-tailed t test was used to calculate P-values. A detailed description of all methods used in this study can be found in *SI Appendix*.

## Acknowledgements

We would like to thank members of the Samuelson laboratory, past and present, for their thoughtful insight and assistance related to this project. We would like to thank the laboratories of Drs. Emily Troemel (UCSD), David Wang (WUSTL), and Gary Ruvkun (MGH/HMS) for reagents. Some strains were provided either by the CGC, which is funded by NIH Office of Research Infrastructure Programs (P40 OD010440), or by NBP-Japan. We thank Dr. Maureen Ferran (RIT) for many discussions and advice. We thank Dr. Mary Wines-Samuelson for critical reading of this manuscript. We would also like to thank members of the Department of Biomedical Genetics (URMC) and the Western New York Worm Group for helpful discussions and feedback, particularly Drs. Doug Portman, Benoit Biteau, Andrew Wojtovich, and Keith Nehrke (URMC). We would like to thank Schrödinger’s cat for staying in the box while this manuscript was in preparation. Research reported in this publication was supported by the National Institute on Aging and the National Institute of General Medical Sciences of the National Institutes of Health under Award Numbers R21AG064519, RF1AG062593 and R21GM148859, respectively. The content is solely the responsibility of the authors and does not necessarily represent the official views of the National Institutes of Health.

