## Supplemental Appendix for "Antiviral defense in aged *Caenorhabditis elegans* declines due to loss of DRH-1/RIG-I deSUMOylation via ULP-4/SENP7"

### Supporting Information Text

#### Extended Materials and Methods

***C. elegans* strains and maintenance.** *C. elegans* were maintained on nematode growth medium (NGM) plates seeded with OP50 *Escherichia coli*. Animals were maintained at 20°C unless otherwise noted. See Table S1 for a list of all *C. elegans* strains used in this study.

**Table S1. List of strains used in this study.**

| Strain name | Genotype | Figures | Source |
|---|---|---|---|
| ERT54 | *jyIs8 [pals-5p::GFP + myo-2p::mCherry]* | 1,S1,S2,2,S3,3,6,7,S9 | CGC |
| AVS876 | *ulp-4(tm3688)/mIn1 [mIs14 dpy-10(e128)]; jyIs8 [pals-5p::GFP + myo-2p::mCherry]* | 1,2,S3 | This study |
| AVS877 | *ulp-4(tm3688)/mIn1 [mIs14 dpy-10(e128)] II; jyIs8 [pals-5p::GFP + myo-2p::mCherry]; artEx95[ges-1p::ulp-4 + myo-2p::mCherry]* | 2,S3 | This study |
| AVS878 | *ulp-4(tm3688)/mIn1 [mIs14 dpy-10(e128)] II; jyIs8[pals-5p::GFP + myo-2p::mCherry]; artEx96[myo-3p::ulp-4 + myo-2p::mCherry]* | S3 | This study |
| AVS912 | *ulp-4(tm3688)/mIn1 [mIs14 dpy-10(e128)] II; jyIs8[pals-5p::GFP + myo-2p::mCherry]; artEx104[dpy-7p::ulp-4 + myo-2p::mCherry]* | S3 | This study |
| AVS911 | *ulp-4(tm3688)/mIn1 [mIs14 dpy-10(e128)] II; jyIs8[pals-5p::GFP + myo-2p::mCherry]; artEx103[rab-3p::ulp-4 + myo-2p::mCherry]* | S3 | This study |
| VP303 | *rde-1(ne219) V; kbIs7 [nhx-2p::rde-1 + rol-6(su1006)]* | 2 | CGC |
| AVS881 | *gei-17(tm2723)/ hT2 [bli-4(e937) let-?(q782) qIs48] I; jyIs8 [pals-5p::GFP + myo-2p::mCherry]* | S1 | This study |
| AVS882 | *drh-1(ok3495);jyIs8[pals-5p::GFP+ myo-2p::mCherry]* | 3,6 | This study |
| GR3325 | *mgTi54[rpl-28p::mScarlet::drh-1]* | 3 | Ruvkun laboratory |
| AVS928 | *artIs14[rpl-28p::mScarlet::drh-1+ myo-2p::gfp]* | 4,S4,S5,7,S8,S9 | This study |
| AVS929 | *ulp-4(tm3688)/mIn1 [mIs14 dpy-10(e128)]; artIs14[rpl-28p::mScarlet::drh-1+ myo-2p::gfp]* | 4 | This study |
| AVS932 | *ulp-4(tm3688)/mIn1 [mIs14 dpy-10(e128)]; artIs14[rpl-28p::mScarlet::drh-1+ myo-2p::gfp]; artEx95[ges-1p::ulp-4 + myo-2p::mCherry]* | 4,S5 | This study |
| AVS1036 | *artIs14[rpl-28p::mScarlet::drh-1+ myo-2p::gfp]; zcIs17[ges-1p::GFP(mit)].* | 4 | This study |
| AVS1037 | *artIs14[rpl-28p::mScarlet::drh-1+ myo-2p::gfp]; [smo-1p::8XHIS::GFP::smo-1]* | 4 | This study |
| AVS1038 | *drh-1(ok3495);jyIs8[pals-5p::GFP+ myo-2p::mCherry]; artEx106 [rpl-28p::mScarlet::drh-1+ myo-2p::gfp]* | 5,S6,7 | This study |
| AVS1039 | *drh-1(ok3495);jyIs8[pals-5p::GFP+ myo-2p::mCherry]; artEx107[rpl-28p::mScarlet::drh-1(k647R+K731R)+ myo-2p::gfp]* | 5,S6,7 | This study |
| AVS1040 | *artEx106[rpl-28p::mScarlet::drh-1+myo-2p::gfp]; zcIs17 [ges-1p::GFP(mit)]* | S7 | This study |
| AVS1041 | *artEx107[rpl-28p::mScarlet::drh-1(k647R+K731R)+ myo-2p::gfp]; zcIs17 [ges-1p::GFP(mit)]* | S7 | This study |
| ERT60 | *jyIs13 [act-5p::GFP::ACT-5 + rol-6(su1006)] II* | 6 | CGC |
| AVS914 | *drh-1(ok3495);jyIs8[pals-5p::GFP+ myo-2p::mCherry]; artIs11 [ges-1p::6XHIS::drh-1+ myo-2p::gfp]* | 7 | This study |
| AVS926 | *drh-1(ok3495);jyIs8[pals-5p::GFP+ myo-2p::mCherry]; artIs12[ges-1p::6XHIS::drh-1(k647R+K731R)+myo-2p::gfp]* | 7 | This study |

#### Synchronization of *C. elegans*

To age-synchronize animals, gravid adult *C. elegans* were washed off plates with M9 buffer and transferred into a 5 mL conical tube. Clear pellets were resuspended in 2 mL of M9 and 1 mL of bleaching solution (600 $\mu$L of sodium hypochlorite solution and 160 $\mu$L of 5 M NaOH and 240 $\mu$l of H2O). Released embryos were washed two times with 5 mL of M9 and resuspended in a final volume of 3 mL of M9. Embryos were incubated at 20 $^{o}$C with continual rotation for 20-24 h to hatch synchronized L1s.

#### *C. elegans* RNAi

RNA interference was achieved by bacterial feeding using clones from the MRC RNAi library, after verifying sequence identity. All RNAi bacterial were not diluted, with one exception. *ubq-2(RNAi)* can cause developmental arrest, therefore bacteria were diluted 10-fold with either empty vector RNAi or RNAi indicated within the text. Synchronized L1 were fed with empty vector at 20°C for 20 hours then transferred to diluted *ubq-2(RNAi)* to prevent developmental arrest.

**Orsay virus infection**

For all strains, except those with *ulp-4(tm3688)*, synchronized L1 larvae were put onto 6 cm RNAi plates and incubated at 16°C until the L4 stage, then a 200 uL of 1:1 dilution of Orsay Virus was top added onto the plates, which were dried at room temperature. Infected worms were incubated at 20°C until collection. For AVS876-878, and AVS911-912, synchronized L1 larvae on 10 cm empty vector RNAi plates were incubated at 16°C, then at L4 approximately100 *ulp-4 (tm3688)* homozygotes, with or without transgenes, were moved to 6 cm plates for infection.

**Bortezomib treatment**

Synchronized L1 were plated onto 6 cm HT115 empty vector plates and incubated at 16°C until L4. A 10 μM stock solution of bortezomib (Selleck Chemicals), resuspended in dimethyl sulfoxide (DMSO), was mixed with M9 and top plated onto plates for a final concentration of 2.5 uM bortezomib per plate. Control plates were top plated with an equal amount of DMSO in M9 buffer. Plates were then incubated at 20°C for 24 h prior to collection in TRIZOL.

**Prolonged heat stress**

Synchronized L1 were plated onto 6 cm HT115 empty vector plates and incubated at 16°C until L4. Experimental plates were then incubated at 28°C for 24 h, while control plates were incubated at 20°C. Animals were collected in TRIZOL after 24 h.

**Molecular Biology and Transgenesis**

To get a fusion construct containing the *ulp-4* coding sequence, the GFP fragment of pPD95.75 vector was replaced by 4736 bp of the *ulp-4* genomic sequence fragment starting from the start codon of the *ulp-4* open reading frame. For intestinal expression, *ges-1p::ulp-4* was generated by PCR-cloning the 2073 bp *ges-1* promoter fragment upstream of *ulp-4* coding sequence. For epidermal expression, *ges-1p* was replaced by 803 bp of the *dpy-7* promoter. For expression in muscle, *ges-1p* was replaced by 2681 bp of the *myo-3* promoter. For neuronal expression, *ges-1p* was replaced by 1601 bp of the *rab-3* promoter. Tissue specific ULP-4 expression plasmids were then injected into AVS876 at 25 ng/μL; *myo-2P::mCherry* was used as co-injection marker.

The *rpl-28p::mScarlet::drh-1* plasmid was a gift from the laboratory of Gary Ruvkun (MGH/HMS). NBE Q5® Site-Directed Mutagenesis Kit (E0554) was used to generate the *rpl-28p::mScarlet::drh-1(K647A+K731A)* construct. *rpl-28p::mScarlet::drh-1* and *rpl-28p::mScarlet::drh-1(K647A+K731A)* were injected into AVS882 at 25 ng/μL; *myo-2p::GFP* was used as co-injection marker.

To construct *ges-1p::6xHIS::drh-1*, the *ulp-4* genomic sequence of *ges-1p::ulp-4* was replaced by the *drh-1* genomic sequence with 6XHIS tagged at the N terminus. The NBE Q5® Site-Directed Mutagenesis Kit (E0554) was used to generate *ges-1p::6xHIS::drh-1(K647A+K731A)*; this construct or *ges-1P::6xHIS::drh-1* were injected into AVS882 at 10 ng/μL; *myo-2p::GFP* was used as co-injection marker.

Constructs were generated with the NEBuilder HiFi DNA Assembly Cloning Kit (NEB #E5520). All fragments obtained by PCR amplification were confirmed by sequencing.

**TMP/UV Integration**

L2-L4 stage transgenic animals expressing an extrachromosomal array were collected in M9, washed, and resuspended in 200 μL of M9. TMP solution was added to obtain a final concentration 100 μg/mL and animals were incubated at RT for 30 minutes in the dark. Animals were then transferred to an unseeded 100 mm plate and exposed to 350 μJ (x100) long wave UV in a Stratalinker 1800. F2 animals with a 90-100% expression of a transgene in F3 progeny were putative homozygotic integrated transgenic lines, which was confirmed and then 6x backcrossed to N2.

**RNA isolation and RT-qPCR**

Animals were isolated with M9, washed until clear, 200 μL TRIZOL was added to the pellet and frozen at -80°C. RNA was extracted using TRIZOL reagent according to the manufacturer's instructions. cDNA was prepared from total RNA using iScript (Bio-Rad) cDNA synthesis kits. RT-qPCR was performed using PerfeCTa® SYBR® Green FastMix®, Low ROX™ (VWR) on a Quantum studio 5 system. Each replicate was measured in triplicate. All gene expression was normalized to *snb-1* and *cdc-42* expression. RT-qPCR primer sequences are listed in Table S2.

**Table S2. List of QPCR primers used in this study**

| Primer Name | Sequence |
|---|---|
| cdc-42-F | CGACAATTACGCCGTCACAG |
| cdc-42-R | AAACACGTCGGTCTGTGGAT |
| snb-1-F | CCGGATAAGACCATCTTGACG |
| snb-1-R | GACGACTTCATCAACCTGAGC |
| pals-5-F | GGACTATGTGAGCATATTGTTTCC |
| pals-5-R | AGCGTTTTTCTGCACTGGTT |
| RNA1-F | ACCTCACAACTGCCATCTACA |
| RNA1-R | GACGCTTCCAAGATTGGTATTGGT |
| drh-1-F | GCGATTTTCACTACATTCTCGA |
| drh-1-R | TTCTTCGTTGCGCTCTATTTC |
| ulp-4-F | CGGAACTCATCTAGATGGTTCTATC |
| ulp-4-R | CCGCTTCTGATTTGTCCATT |
| sdz-6-F | ACAATCGGGCGTTCAATTC |
| sdz-6-R | TCTGATAGCTGGCTGAGTGG |

**Fluorescence Microscopy of *C. elegans***

Animals were added to a 10 mM levamisole droplet on 2% agarose pads. Images were captured with the Leica DMi8 confocal microscope or Zeiss Axio Imager.M2m and processed with Fiji.

**Immunoblotting**

Animals raised under each described condition isolated with M9 and washed until the supernatant was clear. 2X SDS Laemmli buffer (4% SDS, 20% glycerol, 10% 2-mercaptoethanol, 0.004% bromophenol blue, 0.125M tris-HCl, pH 6.8) was used to resuspend the pellet. Samples were boiled at 97°C for 10 min. Normalized protein levels were loaded onto 7% SDS-PAGE and transferred onto Nitrocellulose membranes (Bio-Rad). After blocking with 5% non-fat milk, the membrane was probed with the designated primary and secondary antibodies (rabbit polyclonal anti-RFP, Rockland # 600-401-379; rabbit polyclonal anti-actin, CST #4967; Goat anti-Rabbit IgG (H+L) Secondary Antibody, Thermo Fisher # 31460), and developed with SuperSignal™ West Atto Ultimate Sensitivity Substrate (Thermo Fisher # A38555), and visualized by Bio-Rad ChemiDoc Imaging System. Analysis was performed using Fiji.

**Quantification and statistical analysis**

Relative *pals-5p::GFP* and *mScarlet::DRH-1* expression was analyzed by Fiji. Background mean fluorescence intensity was subtracted from each animal.

Immunoblot quantification was analyzed by Fiji.

All statistical analysis was performed with GraphPad Prism. A two-tailed t test was used to calculate P-values; *P < 0.05. **P < 0.01, ***P < 0.001 and ****P < 0.0001.

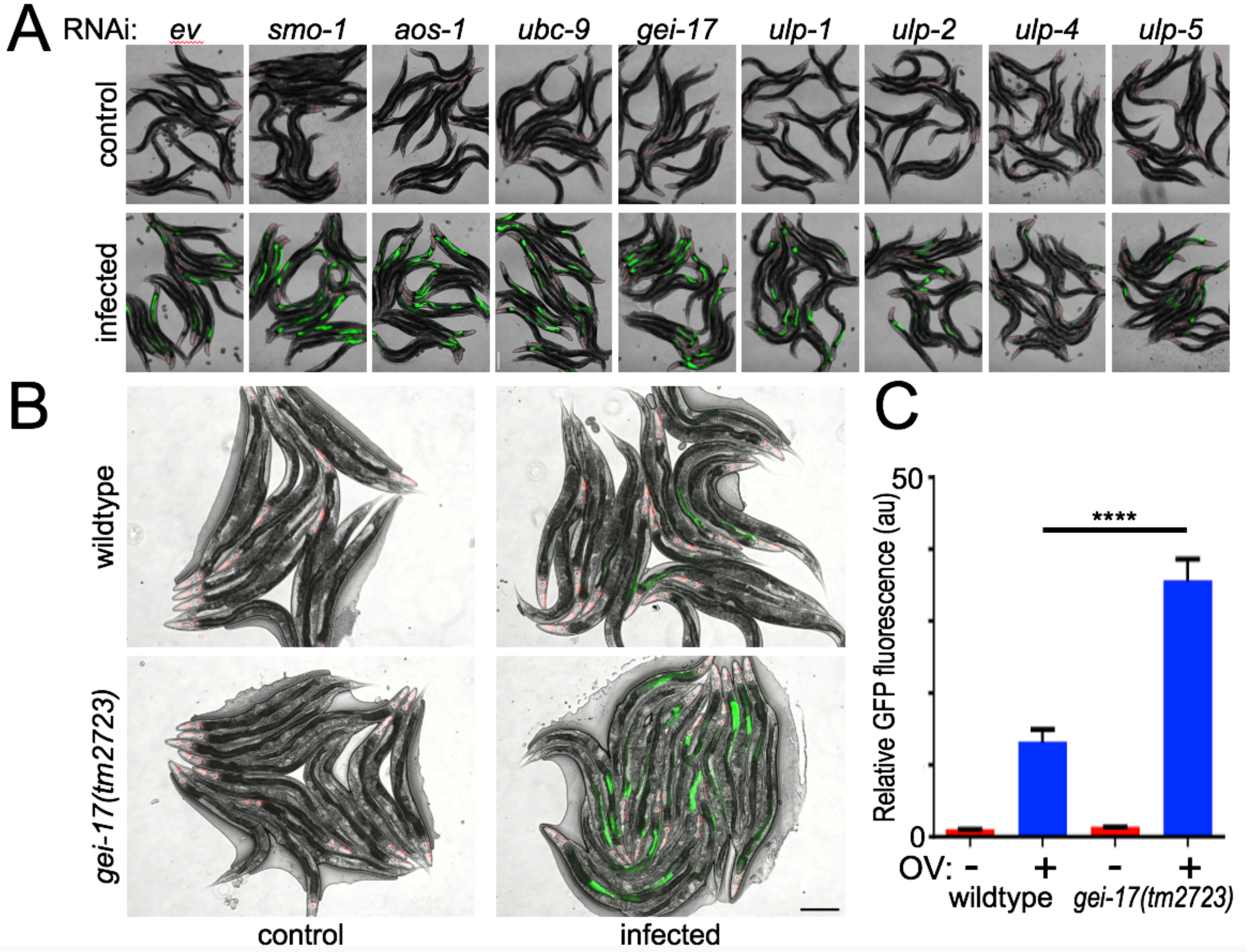


***SI Appendix*, Fig. 1. Inhibition of SUMOylation amplifies IPR induction after viral infection.** (*A*) Representative images of *pals-5p::GFP* expression after RNAi of the indicated component of the SUMOylation machinery +/- viral infection. Quantification of GFP fluorescence is in Fig. 1*B.* (*B*) Representative images of *pals-5p::GFP* expression in infected wildtype and *gei-17(tm2723)* null mutant animals. Scale bar = 200 µm. (*C*) Quantification of GFP fluorescence intensity of samples in (*B*). In all cases, values are the mean of 60 animals across three independent trials; error bars represent the SEM. A two-tailed t test was used to calculate P-values; ****P < 0.0001.

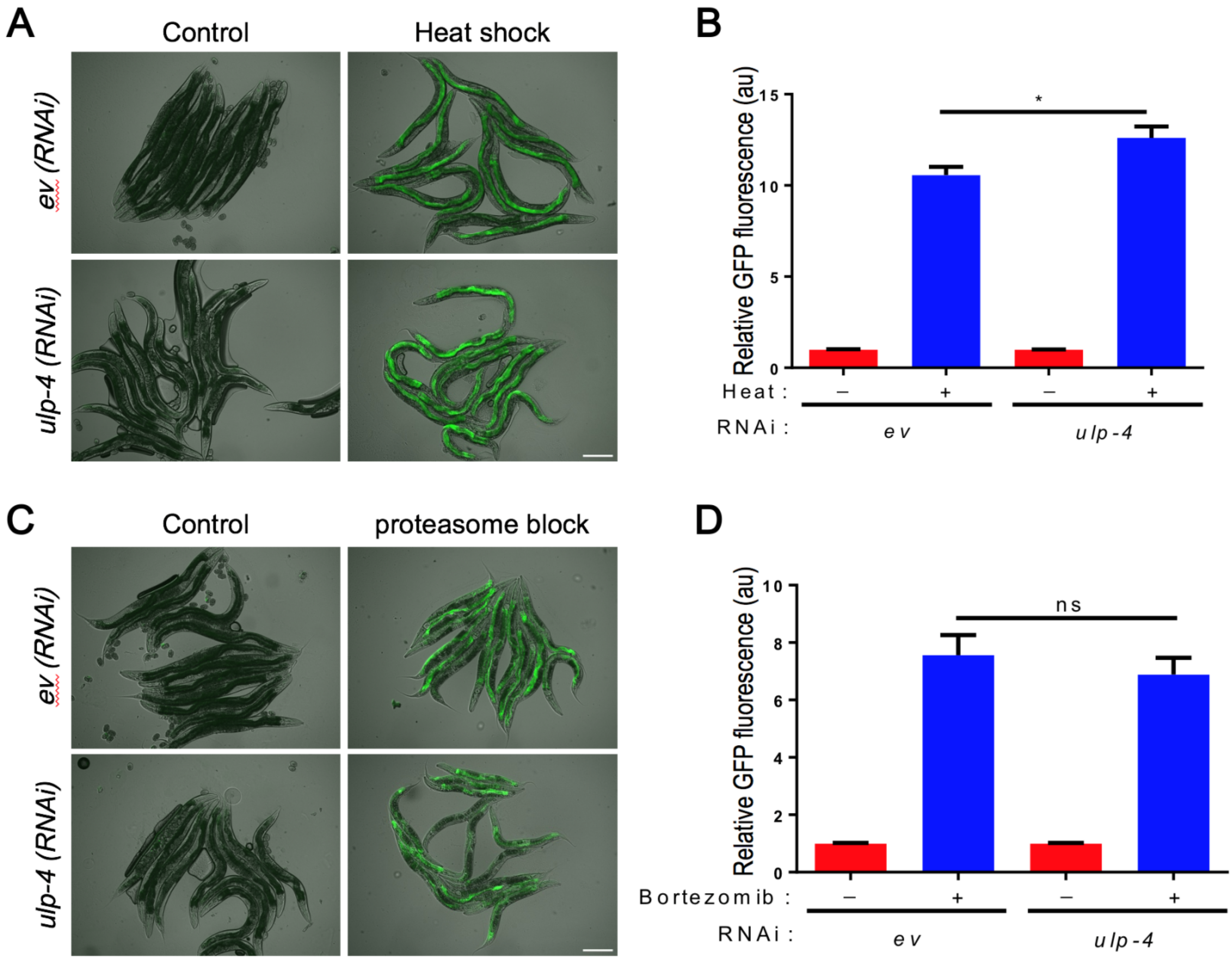


***SI Appendix*, Fig. 2. ULP-4 is not required for induction of the IPR by non-viral triggers.** (*A*) Representative images of *pals-5p::GFP* expression after empty vector or *ulp-4(RNAi)* treatment, followed by 28ºC heat stress. (*B*) Quantification of GFP fluorescence in (*A*). (*C*) Representative images of *pals-5p::GFP* expression after empty vector or *ulp-4(RNAi)* treatment, followed by bortezomib treatment. (*D*) Quantification of GFP fluorescence in (*C*). Scale bar = 200 µm. In all cases, values are the mean of 60 animals across three independent trials; error bars are the SEM. A two-tailed t test was used to calculate P-values; *$P < 0.05$, ns = no significant difference.

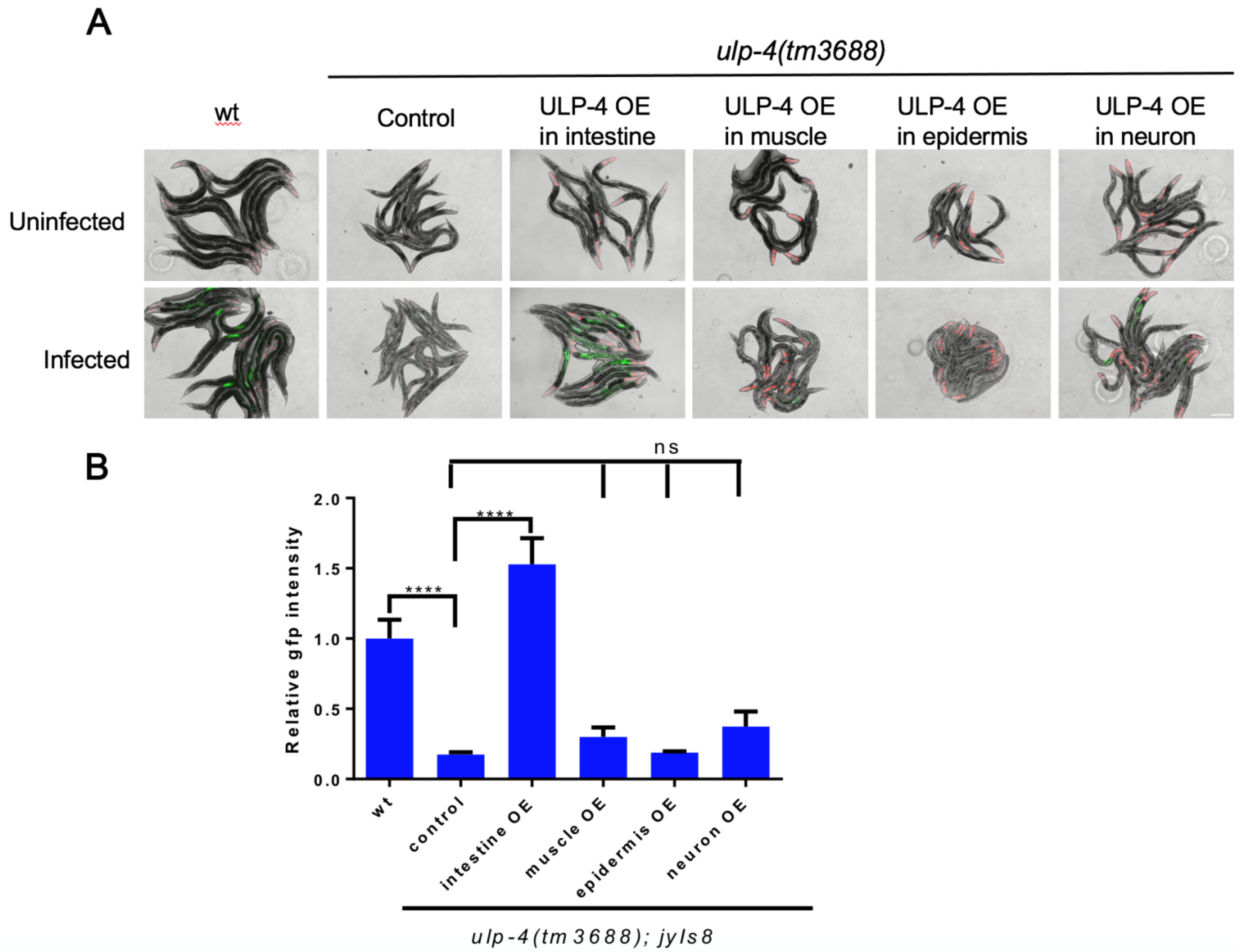


***SI Appendix*, Fig. 3. ULP-4 does not act outside of the intestine to regulate IPR following viral infection.** *(A)* Representative images of *pals-5p::GFP* expression +/- viral infection of animals that were: wildtype, *ulp-4(tm3688)*, *ulp-4(tm3688);artEx95(ges-1p::ULP-4), ulp-4(tm3688);artEx96(myo-3p::ULP-4), ulp-4(tm3688);artEx104(dpy-7p::ULP-4),* or *ulp-4(tm3688);artEx103(rab-3p::ULP-4)*, which restored ULP-4 expression within the intestine, body wall muscle, epidermis, or nervous system, respectively. Scale bar = 200 µm. (*B*) RT-qPCR analysis of endogenous *pals-5* mRNA of samples in (*A*). In all cases, values are the mean of 60 animals across three independent trials; error bars are SEM. A two-tailed t test was used to calculate P-values; ****P < 0.0001, ns = no significant difference.

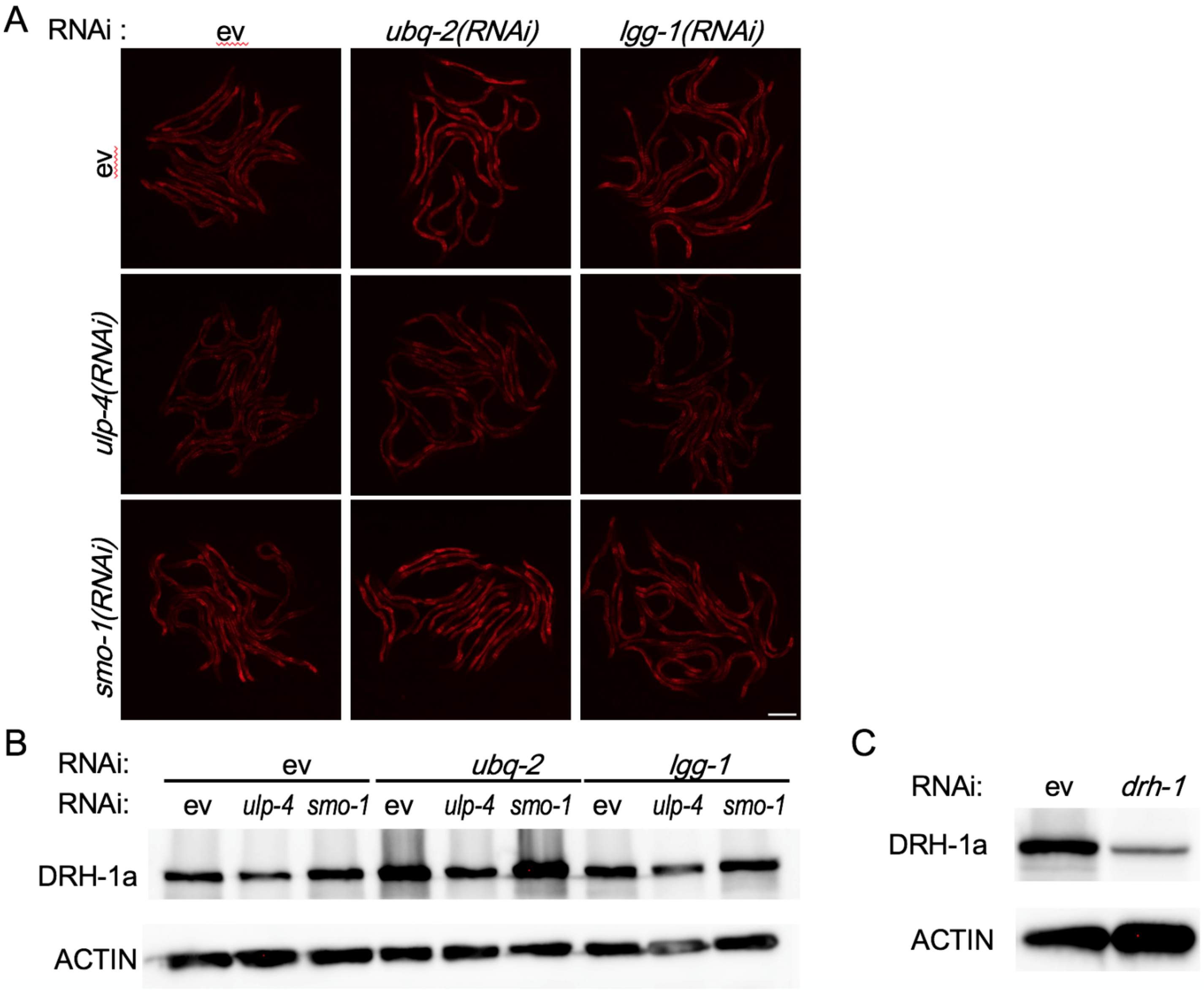


***SI Appendix*, Fig. 4. DRH-1 is degraded through proteasome.** (*A*) Representative images of *mScarlet::DRH-1* expression after RNAi treatment with empty vector, *ubq-2*, *lgg-1*, *ulp-4*, *ulp-4+ubq-2*, *ulp-4+lgg-1*, *smo-1*, *smo-1+ubq-2,* or *smo-1+lgg-1,* respectively. Scale bar = 200 µm. (*B*) Representative immunoblot of DRH-1 levels. (*C*) Representative immunoblot of DRH-1 levels after empty vector or *drh-1(RNAi)*.

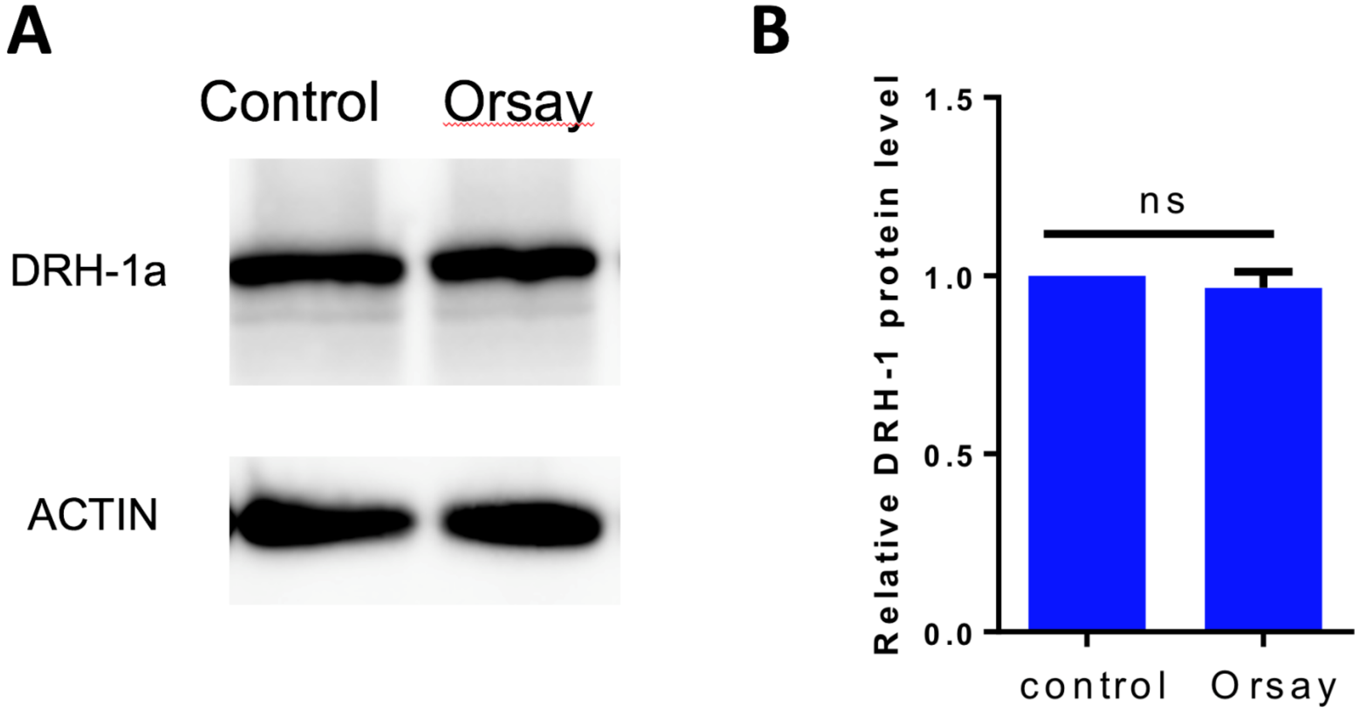


***SI Appendix*, Fig. 5. Viral infection does not change DRH-1 protein levels.** (*A*) Representative immunoblot of mScarlet::DRH-1 in animals +/- viral treatment. (*B*) Quantification of DRH-1 levels. Values are the mean from three independent experimental replicates; error bars are the SEM. A two-tailed t test was used to calculate P-values; ns = no significant difference.

**A**

| position | sequence | SUMOplot | JASSA | GPS-SUMO | MusiteDeep |
|---|---|---|---|---|---|
| K647 | VALNY **L<u>K</u>DE** MEYRT | 0.91 | LOW | 0.735 | 0.764 |
| K731 | LLMLG **I<u>K</u>SE** WMSGL | 0.94 | HIGH | 0.826 | 0.926 |

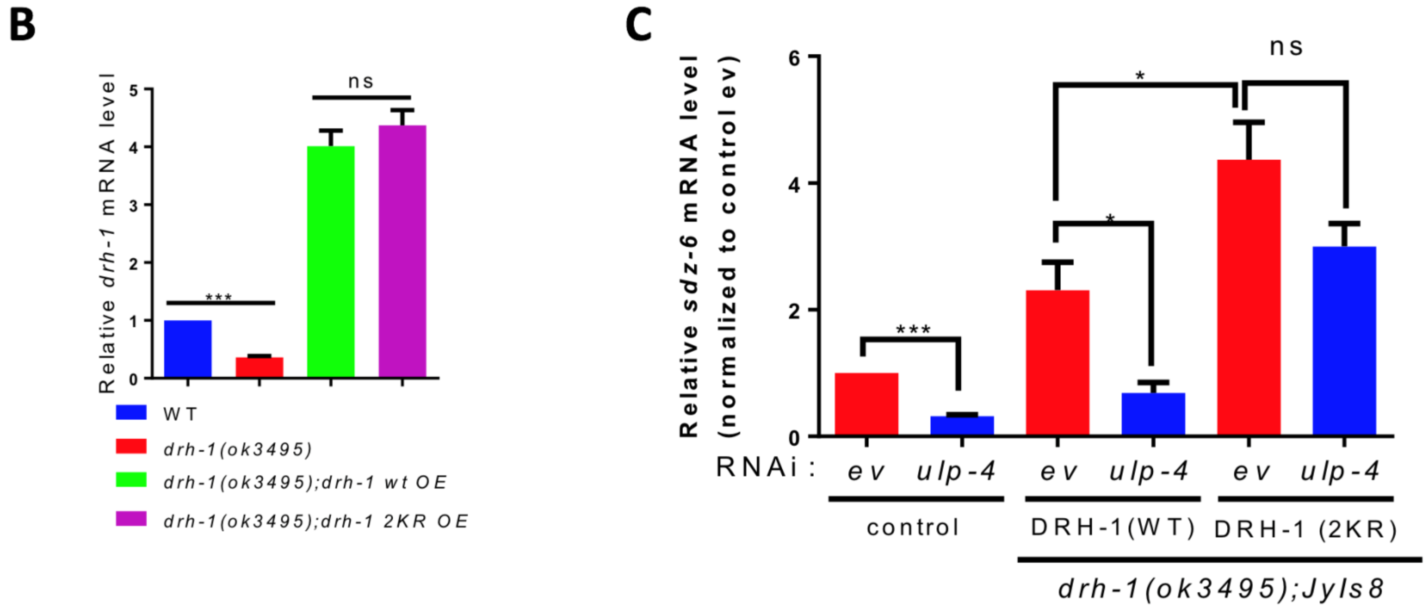


***SI Appendix*, Fig. 6. K647/K731 are the predicted target residues for DRH-1 SUMOylation.** (*A*) SUMOylation likelihood scores for DRH-1 lysines 647 and 731 as predicted by four independent computational models. The detailed likelihood for all 60 DRH-1 lysines are listed in *Dataset S1*. (*B*) RT-qPCR analysis of endogenous *drh-1* in wildtype, *drh-1(ok3495)* null mutant, and *drh-1(ok3495)* null mutant animals with restored expression throughout the soma (*rpl-28p*) of either wildtype DRH-1 (*mScarlet::DRH-1(wt)*) or nonSUMOylatable DRH-1 (*mScarlet::DRH-1(2KR)*). (*C*) RT-qPCR analysis of endogenous *sdz-6* under the indicated conditions. In all cases, values are the mean of three independent trials; error bars are the SEM. A two-tailed t test was used to calculate P-values; *P < 0.05,***P < 0.001, ns = no significant difference.

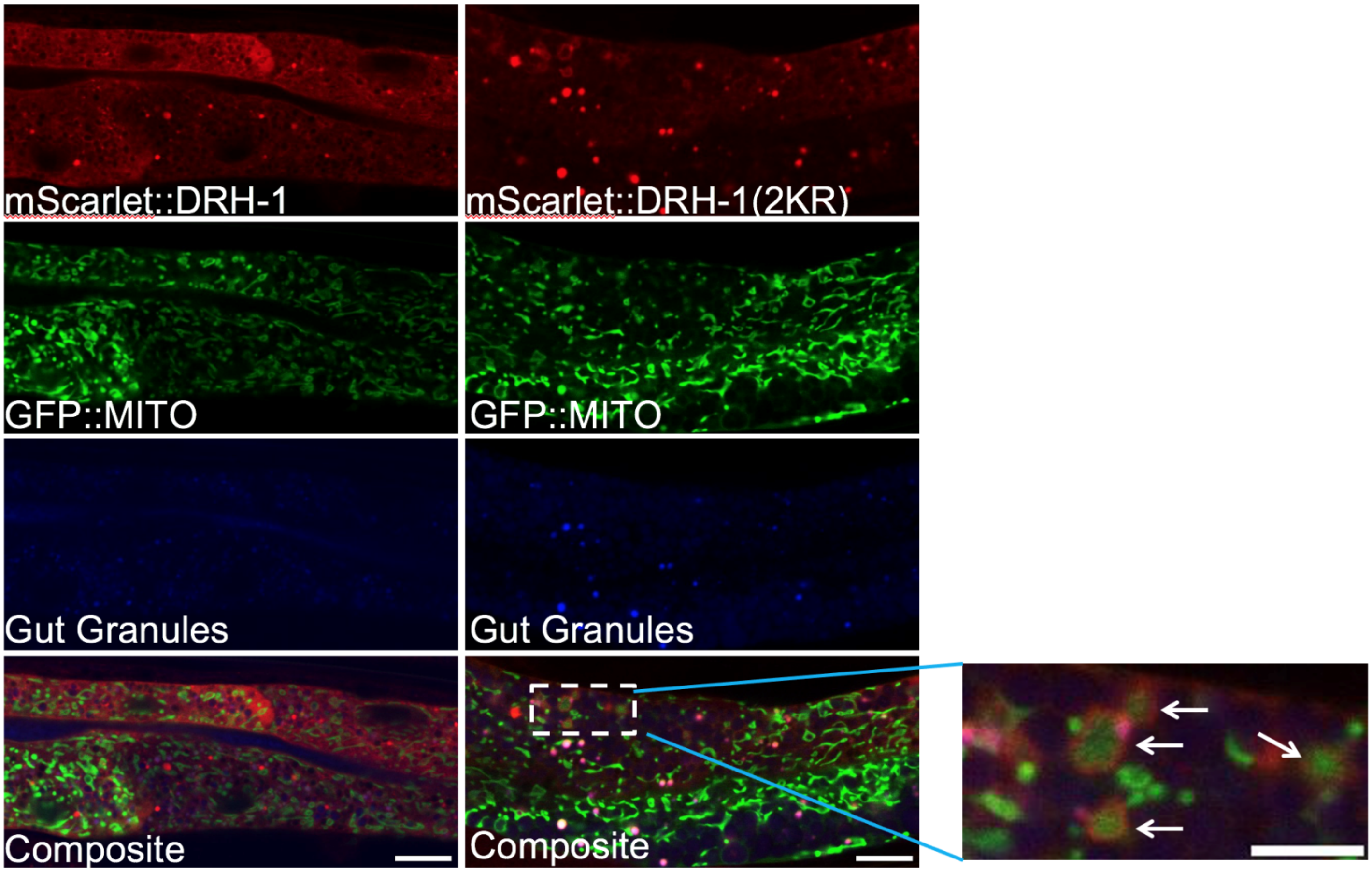


***SI Appendix*, Fig. 7. nonSUMOylated DRH-1 co-localizes with mitochondria.** Representative images of uninfected animals expressing *mScarlet::DRH-1(WT); GFP::mito* or *mScarlet::DRH-1(2KR); GFP::mito* respectively, lysosomal related organelles/gut granules are indicated (405-nm blue channel). Scale bar = 2 μm. Inset scale bar = 5 μm

.

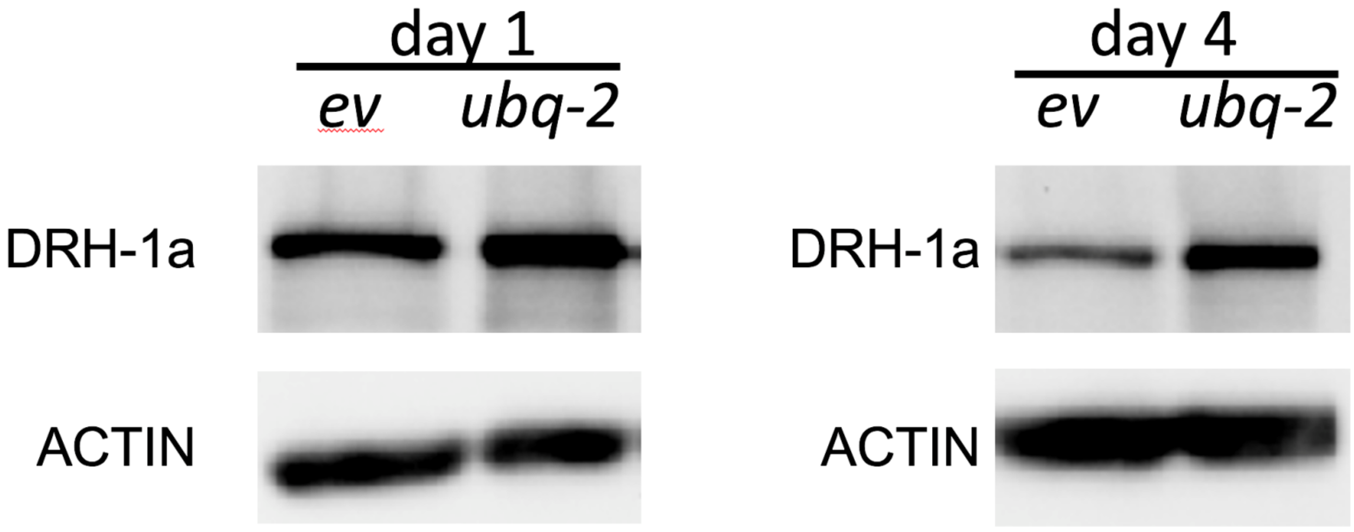


***SI Appendix*, Fig. 8. Aging results in increased proteosomal degradation of DRH-1.** Representative immunoblot of *mScarlet::DRH-1* expression at the indicated ages after either empty vector or *ubq-2(RNAi)* treatment.

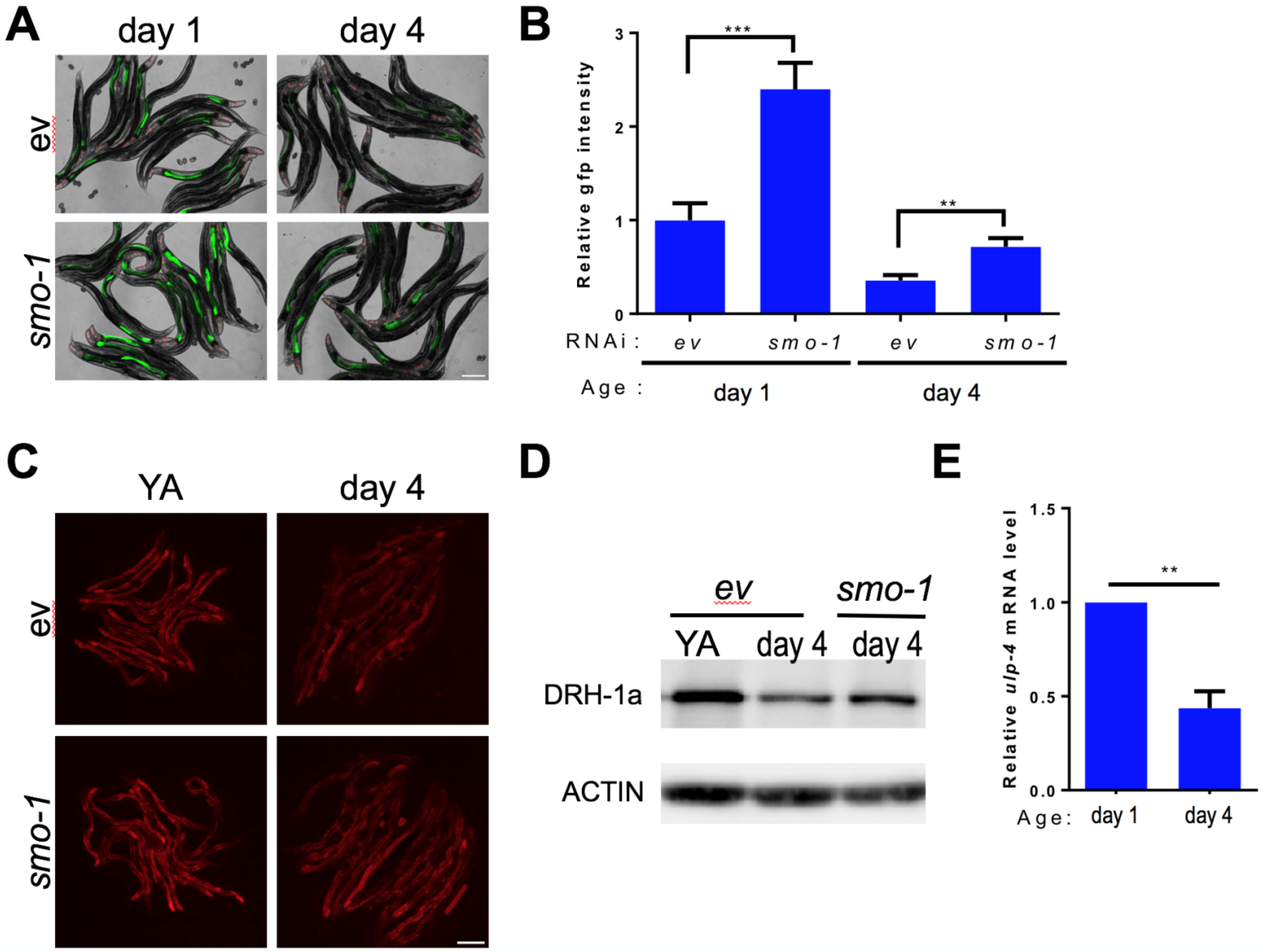


***SI Appendix*, Fig. 9. Inhibition of SUMOylation partially rescued the declined inducibility of the IPR in older animals.** (*A*) Representative images of *pals-5p::GFP* expression +/- viral infection. Feeding based RNAi was initiated at L1, viral infection was initiated at either L4 or day 3 of adulthood, and images were obtained 24 hours later. Scale bar = 200 μm. (*B*) Quantification of GFP fluorescence from (*A*). Values are the relative mean of 60 animals across three independent trials; error bars are the SEM. A two-tailed t test was used to calculate P-values; $^{**}P < 0.01$, $^{***}P < 0.001$. (*C*) Representative images of *mScarlet::DRH-1* expression after either empty vector or *smo-1(RNAi)*. Scale bar = 200 μm. (*D*) Representative immunoblot of samples analogous to (*C*). (*E*) RT-qPCR analysis of endogenous *ulp-4* in either day 1 or day 4 adult, wildtype animals.
